# Human NAIP/NLRC4 and NLRP3 inflammasomes detect *Salmonella* type III secretion system activities to restrict intracellular bacterial replication

**DOI:** 10.1101/2021.06.17.448811

**Authors:** Nawar Naseer, Marisa Egan, Valeria M. Reyes Ruiz, Igor E. Brodsky, Sunny Shin

**Author notes:** Department of Pathology, Microbiology, and Immunology, Vanderbilt University Medical Center, Nashville, TN 37232. These authors contributed equally to the work (Nawar Naseer, Marisa Egan, Valeria M. Reyes Ruiz).

## Abstract

*Salmonella enterica* serovar Typhimurium is a Gram-negative pathogen that uses two distinct type III secretion systems (T3SSs), termed *Salmonella* pathogenicity island (SPI)-1 and SPI-2, to deliver virulence factors into the host cell. The SPI-1 T3SS enables *Salmonella* to invade host cells, while the SPI-2 T3SS facilitates *Salmonella*’s intracellular survival. In mice, a family of cytosolic immune sensors, including NAIP1, NAIP2, and NAIP5/6, recognizes the SPI-1 T3SS needle, inner rod, and flagellin proteins, respectively. Ligand recognition triggers assembly of the NAIP/NLRC4 inflammasome, which mediates caspase-1 activation, IL-1 family cytokine secretion, and pyroptosis of infected cells. In contrast to mice, humans encode a single NAIP that broadly recognizes all three ligands. The role of NAIP/NLRC4 or other inflammasomes during *Salmonella* infection of human macrophages is unclear. We find that although the NAIP/NLRC4 inflammasome is essential for detecting SPI-1 T3SS ligands in human macrophages, it is partially required for responses to infection, as *Salmonella* also activated the NLRP3 and CASP4/5 inflammasomes. Importantly, we demonstrate that combinatorial NAIP/NLRC4 and NLRP3 inflammasome activation restricts *Salmonella* replication in human macrophages. In contrast to SPI-1, the SPI-2 T3SS inner rod is not sensed by human or murine NAIPs, which is thought to allow *Salmonella* to evade host recognition and replicate intracellularly. Intriguingly, we find that human NAIP detects the SPI-2 T3SS needle protein. Critically, in the absence of both flagellin and the SPI-1 T3SS, the NAIP/NLRC4 inflammasome still restricted intracellular *Salmonella* replication. These findings reveal that recognition of *Salmonella* SPI-1 and SPI-2 T3SSs and engagement of both the NAIP/NLRC4 and NLRP3 inflammasomes control *Salmonella* infection in human macrophages.

**Author summary:** *Salmonella enterica* serovar Typhimurium is a gastrointestinal bacterial pathogen that causes diarrheal disease and is a major cause of morbidity and mortality worldwide. *Salmonella* uses molecular syringe-like machines called type III secretion systems (T3SSs) to inject virulence factors into host cells. These T3SSs enable *Salmonella* to infect and survive within host cells such as macrophages. However, host cells contain a family of cytosolic immune receptors, termed NAIPs, that recognize T3SS and flagellin components. Upon detecting these components, NAIPs recruit the adaptor protein NLRC4 to form signaling complexes called inflammasomes. Inflammasomes activate host proteases called caspases that mount robust immune responses against the invading pathogen. While mice encode multiple NAIPs that have been extensively studied, much remains unknown about how the single human NAIP mediates inflammasome responses to *Salmonella* in macrophages. Our study reveals that while NAIP is necessary to detect individual T3SS ligands in human macrophages, it is only partially required for inflammasome responses to *Salmonella* infection. We found that the NLRP3 and CASP4/5 inflammasomes are also activated, and the combination of NAIP- and NLRP3-mediated recognition limits intracellular *Salmonella* replication in human macrophages. Our results demonstrate that human macrophages employ multiple inflammasomes to mount robust host defense against *Salmonella* infection.

## Introduction

*Salmonella enterica* serovar Typhimurium (referred to hereafter as *Salmonella*) is a Gram-negative bacterial pathogen that causes self-limiting gastroenteritis in immune-competent humans. Transmission of *Salmonella* typically occurs upon ingestion of contaminated food or water. Once inside the host, *Salmonella* uses specialized nanomachines known as type III secretion systems (T3SSs) to inject effectors into the host cell cytosol [1]. Subsequently, these effectors remodel host cellular processes to facilitate bacterial colonization. Thus, *Salmonella*’s T3SSs enable the enteric pathogen to successfully colonize the intestinal tract and infect a variety of cell types, including intestinal epithelial cells (IECs) and macrophages [1]. Specifically, *Salmonella* uses its first T3SS, located on *Salmonella* Pathogenicity Island 1 (SPI-1), to invade host cells, and its second T3SS, located on a second pathogenicity island, SPI-2, to persist and replicate within host cells [2–8]. Numerous other Gram-negative bacterial pathogens also use these evolutionarily conserved T3SSs to colonize the host [9]. While T3SSs are required for these bacterial pathogens to cause disease, they also translocate structural components of the T3SS or the flagellar apparatus into the cytosol, thus enabling the host to detect the invading pathogen [10]. Unlike effectors, which display significant diversity across bacterial species, structural components of the T3SS or the flagellar apparatus retain significant structural homology across Gram-negative bacteria [9, 11]. Thus, these ligands serve as ideal targets of host immune sensors.

The mammalian innate immune system is armed with pattern recognition receptors (PRRs) that detect pathogens by recognizing pathogen-associated molecular patterns (PAMPs) [12, 13]. A subfamily of cytosolic PRRs, known as NAIPs (the NLR [nucleotide-binding domain, leucine-rich repeat-containing] family, apoptosis inhibitory proteins), recognize the structurally related SPI-1 T3SS needle protein, SPI-1 T3SS inner rod protein, and flagellin, which are translocated into the host cell cytosol by the SPI-1 T3SS [10,14,15]. Mice have multiple NAIPs, each specific to a particular ligand: NAIP1 recognizes the T3SS needle protein, NAIP2 recognizes the T3SS inner rod protein, and NAIP5 and NAIP6 both recognize flagellin [14–19]. Upon sensing a ligand, NAIPs recruit the adaptor protein NLRC4 (nucleotide-binding domain, leucine-rich repeat-containing family, CARD domain-containing protein 4) to form multimeric signaling complexes called inflammasomes [20–22]. The NAIP/NLRC4 inflammasome then recruits and activates the cysteine protease caspase-1 [23]. Active caspase-1 cleaves downstream substrates, including pro-IL-1 and pro-IL-18, as well as the pore- forming protein gasdermin-D (GSDMD) [24–26]. Cleaved GSDMD creates pores in the host plasma membrane, leading to the release of proinflammatory cytokines and an inflammatory form of cell death known as pyroptosis, which effectively eliminates the infected cell. The NAIP/NLRC4 inflammasome is critical for the control of *Salmonella* infection in mice [27, 28]. However, whether the NAIP/NLRC4 inflammasome recognizes or controls *Salmonella* infection in humans has not been thoroughly investigated.

While mice express several different NAIPs that each respond to a particular ligand, humans only express one functional NAIP [29, 30]. In human macrophages, this single NAIP is sufficient to respond to the cytosolic delivery of bacterial flagellin as well as the SPI-1 T3SS inner rod (PrgJ) and needle (PrgI) proteins [31, 32]. Interestingly, the SPI-2 T3SS inner rod protein (SsaI) fails to induce inflammasome activation in both murine and human macrophages [11, 32], suggesting that the *Salmonella* SPI-2 T3SS evades NAIP detection to enable *Salmonella* replication within macrophages. However, whether the SPI-2 T3SS needle protein (SsaG) is recognized by NAIP or whether NAIP contributes to the restriction of *Salmonella* replication within macrophages is unknown.

In this study, we found that while human macrophages require NAIP and NLRC4 for inflammasome responses to T3SS ligands, NAIP and NLRC4 are only partially required for the inflammasome response during *Salmonella* infection. Rather, we found that *Salmonella* infection of human macrophages also activates both the CASP4/5 inflammasome, which senses cytosolic LPS [33], and the NLRP3 inflammasome. Importantly, both the NAIP/NLRC4 and NLRP3 (NLR pyrin domain-containing protein 3) inflammasomes played a functional role in restricting *Salmonella*’s intracellular replication, indicating that they contribute to host defense in a cell-intrinsic manner, as well as via release of inflammatory mediators. Finally, we found that the NAIP/NLRC4 inflammasome recognizes the SPI-2 T3SS needle protein SsaG, and that SPI-1 T3SS and flagellin-independent, NAIP/NLRC4-dependent recognition of *Salmonella* mediates restriction of bacterial replication within human macrophages. Our findings highlight the multifaceted inflammasome response to *Salmonella* infection in human macrophages, and yield important insight into how human macrophages use inflammasomes to sense and respond to intracellular bacterial pathogens.

## Results

### NAIP and NLRC4 are necessary for inflammasome responses to T3SS ligands in human macrophages

In murine macrophages, multiple NAIPs are required for inflammasome responses to the *Salmonella* SPI-1 T3SS inner rod protein (PrgJ), the SPI-1 T3SS needle protein (PrgI), and flagellin [14–19]. In addition, the murine NAIPs and NLRC4 contribute to the inflammasome response during *in vivo Salmonella* infection [11, 19]. In human macrophages, PrgJ, PrgI, and flagellin all activate the inflammasome, while the *Salmonella* SPI-2 inner rod protein (SsaI) does not [31, 32]. Using siRNA-mediated silencing of *NAIP* in human macrophages, we have previously shown that human NAIP is important for maximal inflammasome responses to PrgJ and flagellin [32]. However, siRNA-mediated knockdown of *NAIP* did not completely abrogate inflammasome activation, either due to incomplete knockdown, or the potential contribution of other inflammasomes. Therefore, it remained unclear whether human NAIP or NLRC4 is absolutely required for inflammasome responses to these bacterial ligands or whether additional host sensors also mediate sensing of these ligands.

To test the requirement of the NAIP/NLRC4 inflammasome in human macrophages, we used the Clustered Regularly Interspersed Palindromic Repeat (CRISPR) system, in conjunction with the RNA-guided exonuclease Cas9, to disrupt the *NAIP* and *NLRC4* genes in the human monocytic cell line, THP-1 (Fig. S1A, S2A). We selected one independent single cell clone of *NAIP^-/-^* THP-1s (*NAIP^-/-^* Clone 12) that exhibited reduced *NAIP* mRNA expression by qRT-PCR compared to WT THP-1s (Fig. S1C). Sequence validation confirmed that this clone contained a deletion of 1 or 2 nucleotides in both *NAIP* alleles, resulting in premature stop codons (Fig. S1B). We selected two independent single cell clones of *NLRC4^-/-^* THP-1s (*NLRC4^-/-^* Clone 4 and Clone 7), both of which showed complete loss of NLRC4 protein expression compared to WT THP-1s (Fig. S2D). Both clones were sequence-validated and both alleles of each clone contained mutations that resulted in premature stop codons (Fig. S2B, S2C). These sequence-validated *NAIP^-/-^* and *NLRC4^-/-^* THP-1 clones were used throughout this study.

To test if NAIP and NLRC4 are necessary for sensing and responding to bacterial T3SS ligands, we compared inflammasome responses in wild type (WT), *NAIP^-/-^*, and *NLRC4^-/-^* THP-1 macrophages to T3SS ligands delivered directly into the host cell cytosol. We used the *Bacillus anthracis* toxin system to deliver these bacterial ligands into the cytosol of THP-1s [34]. This system contains two subunits: a protective antigen (PA) that creates a pore in the host endosomal membrane and a truncated lethal factor (LFn) that is delivered through the PA pore into the cytosol. Our T3SS ligands of interest are fused to the N-terminal domain of the *B. anthracis* LFn. When the LFn is added to eukaryotic cells in conjunction with PA (collectively referred to as Tox), the bacterial ligand is delivered directly into the host cell cytosol. Using this system, we delivered a truncated version of *Legionella* flagellin (FlaTox), the *Salmonella* SPI-1 T3SS inner rod protein (PrgJTox), and the *Burkholderia* T3SS needle protein (YscFTox) into THP-1s. We then measured the release of the inflammasome-dependent IL-1 family cytokines IL-1α, IL-1β, and IL-18 and cell death as markers of inflammasome activation. Cells left untreated (Mock) or treated with the PA alone or the LFn fused to the bacterial ligand alone released negligible levels of IL-1β, IL-18, and IL-1α and exhibited minimal cell death (Fig. 1A, 1C, S3A-C, S4A-C). In agreement with previous findings [32], WT THP-1s treated with both the PA and LFn subunits exhibited robust inflammasome activation, and released substantial levels of IL-1β, IL-18, and IL-1α and exhibited considerable cytotoxicity (Fig. 1A, 1C, S3A-C, S4A-C), indicating that robust inflammasome activation requires cytosolic delivery of the ligands. In contrast, both *NAIP^-/-^* THP-1s and *NLRC4^-/-^* THP-1s released negligible levels of inflammasome-dependent cytokines and did not undergo cell death when treated with FlaTox, PrgJTox, or YscFTox (Fig 1A, 1C, S3A-C, S4A-C). Importantly, the *NAIP^-/-^* and *NLRC4^-/-^* THP-1s released IL-1β at levels comparable to those released by WT THP-1s in response to the NLRP3 stimulus LPS + nigericin (Fig. 1B, 1D), indicating that CRISPR/Cas9 editing was specific to the NAIP/NLRC4 inflammasome pathway [35]. In addition, release of the inflammasome-independent cytokine TNF-α was unaffected in *NAIP^-/-^* or *NLRC4^-/-^* THP- 1s (Fig. S3D, S4D). Consistent with our prior results [32] and in agreement with recent studies [36], these results collectively demonstrate that NAIP and NLRC4 are required for inflammasome activation in response to the T3SS inner rod, T3SS needle, and flagellin proteins in human macrophages.

**Fig 1.**
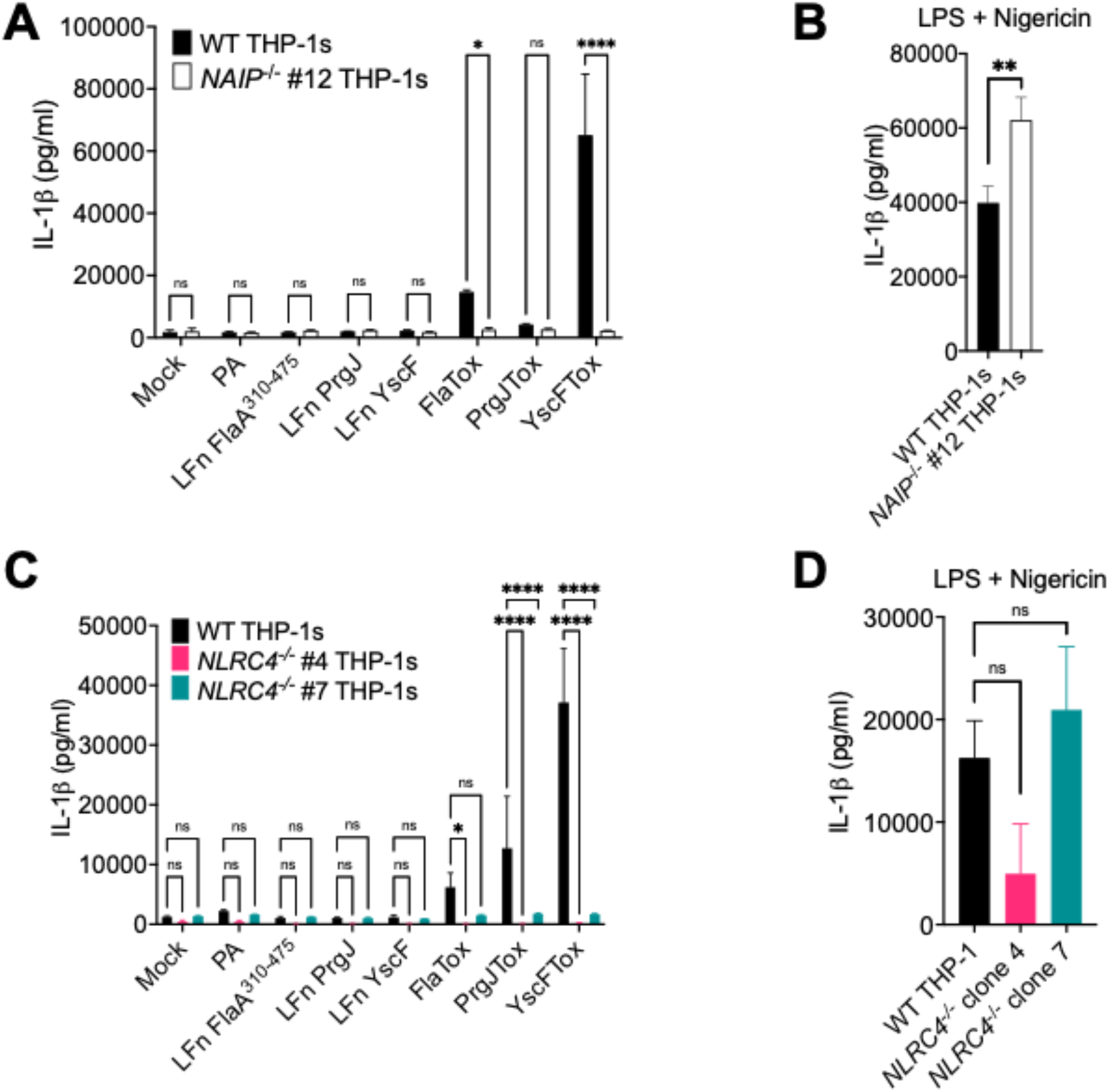
NAIP and NLRC4 are necessary for inflammasome responses to T3SS ligands in human macrophages. WT, *NAIP^-/-^* clone, or two independent clones of *NLRC4^-/-^* THP-1 monocyte-derived macrophages were primed with 100 ng/mL Pam3CSK4 for 16 hours. Cells were then treated with PBS (Mock), PA alone, LFnFlaA^310–475^ alone, LFnPrgJ alone, LFnYscF alone, PA+LFnFlaA^310–475^ (FlaTox), PA+LFnPrgJ (PrgJTox), or PA+LFnYscF (YscFTox) for 6 hours (A, C). As a control, cells were primed with 500 ng/mL LPS for 4 hours and treated with 10 µM nigericin for 6 hours (B, D). Release of IL-1β into the supernatant was measured by ELISA. ns – not significant, \**p* < 0.05, \*\**p* < 0.01, \*\*\*\**p* < 0.0001 by Šídák’s multiple comparisons test (A), or by unpaired t-test (B), or by Dunnett’s multiple comparisons test (C, D). Data shown are representative of at least three independent experiments.

### NAIP and NLRC4 are partially required for inflammasome activation during *Salmonella* infection of human macrophages

Human macrophages undergo SPI-1 T3SS-dependent inflammasome activation during *Salmonella* infection [32]. To test whether this inflammasome activation requires NAIP/NLRC4, we infected WT, *NAIP^-/-^*, or *NLRC4^-/-^* THP-1 macrophages with WT *Salmonella* (WT Stm) or *Salmonella* lacking its SPI-1 T3SS (Δ*sipB* Stm) and assayed for subsequent inflammasome activation (Fig. 2, S5). WT THP-1s infected with WT Stm released high levels of IL-1β, IL-18, and IL-1α and underwent cell death (Fig. 2, S5A-D). This response was dependent on SPI-1 T3SS translocation into host cells, as cells infected with Δ*sipB* Stm, which lack a component of the translocon, failed to undergo robust inflammasome activation (Fig. 2, S5A-D). In *NAIP^-/-^* or *NLRC4^-/-^* THP-1 macrophages infected with WT Stm, we observed a significant decrease but not complete abrogation of secreted IL-1β and IL-18 levels (Fig. 2), whereas levels of IL-1α and cell death were largely unaffected (Fig. S5A–D). WT and *NAIP^-/-^* or *NLRC4^-/-^* THP- 1s released similar levels of the inflammasome-independent cytokine TNF-α (Fig. S5E, S5F). Overall, these data indicate that NAIP and NLRC4 are partially required for inflammasome responses to *Salmonella* infection in human macrophages, in contrast to what we observe with individual T3SS ligand delivery (Fig. 1, S3, S4), where NAIP/NLRC4 is absolutely required for inflammasome activation. Thus, our data indicate that in addition to the NAIP/NLRC4 inflammasome, *Salmonella* also induces a NAIP/NLRC4-independent inflammasome response, in agreement with a recent study [36].

**Fig 2.**
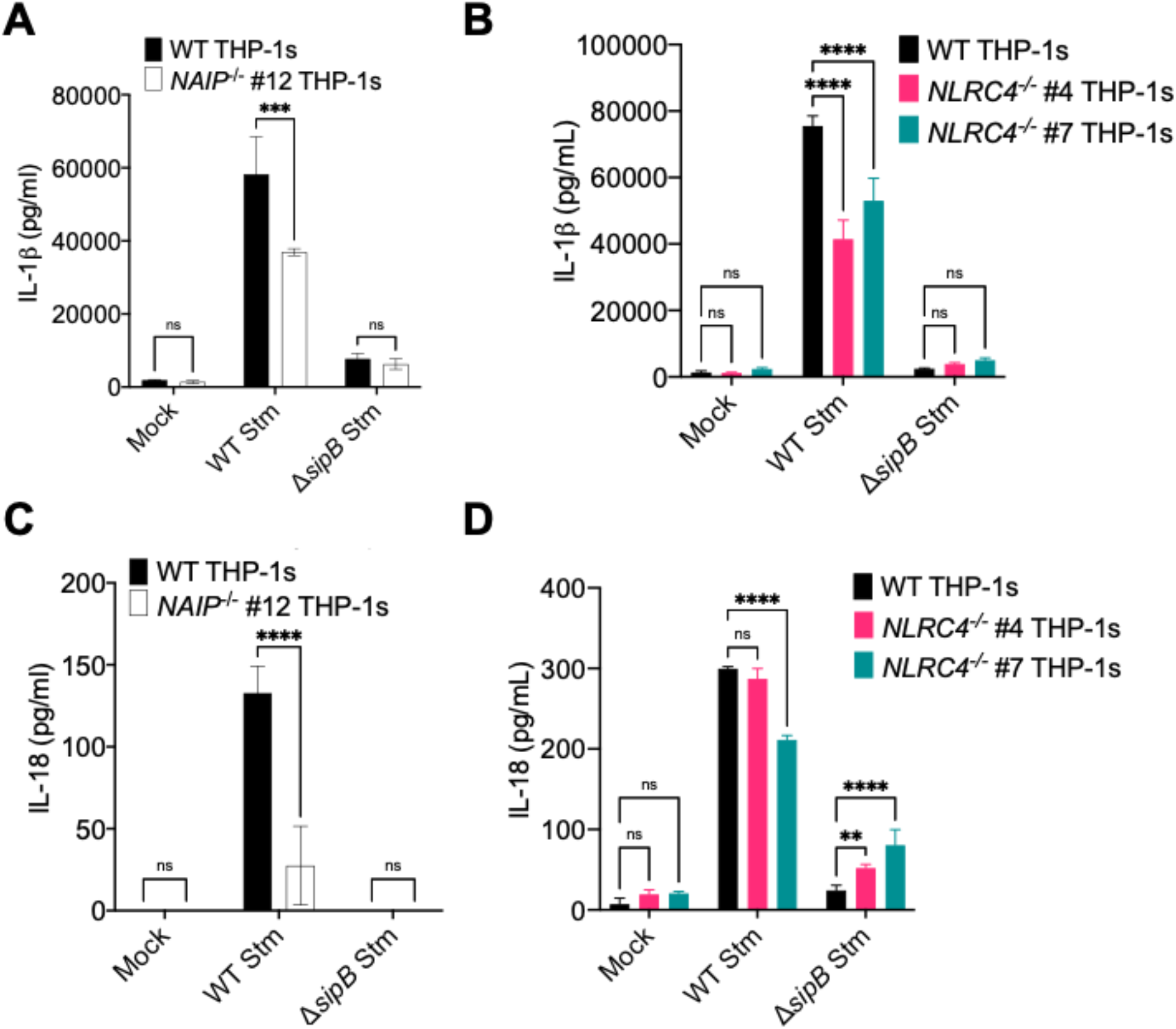
NAIP and NLRC4 are partially required for inflammasome activation during *Salmonella* infection in human macrophages. WT, *NAIP^-/-^* clone, or two independent clones of *NLRC4^-/-^* THP-1 monocyte-derived macrophages were primed with 100 ng/mL Pam3CSK4 for 16 hours. Cells were then infected with PBS (Mock), WT *S*. Typhimurium, or Δ*sipB S*. Typhimurium for 6 hours. Release of IL-1β and IL-18 into the supernatant were measured by ELISA. ns – not significant, \*\*\**p* < 0.001, \*\*\*\**p* < 0.0001 by Šídák’s multiple comparisons test (A, C) or Dunnett’s multiple comparisons test (B, D). Data shown are representative of at least three independent experiments.

### *Salmonella* induces NAIP/NLRC4- and NLRP3-dependent inflammasome activation in human macrophages

In murine macrophages, *Salmonella* infection activates both the NAIP/NLRC4 and NLRP3 inflammasomes [37]. The NAIP/NLRC4 inflammasome is important for early responses to *Salmonella* in the setting of SPI-1 activation, while the NLRP3 inflammasome is important at later timepoints following bacterial replication [38]. In human THP-1s, *Salmonella* infection triggers recruitment of both NLRC4 and NLRP3 to the same macromolecular complex [38]. The NLRP3 inflammasome can be activated by diverse stimuli during bacterial infection, such as potassium efflux [39]. To determine if the NAIP/NLRC4-independent inflammasome response we observed in our *Salmonella*- infected human macrophages is NLRP3-dependent, we infected WT, *NAIP^-/-^*, or *NLRC4^-/-^* THP-1s with *Salmonella* in the presence of MCC950, a potent chemical inhibitor of the NLRP3 inflammasome [40], or the vehicle control DMSO. We subsequently assayed for inflammasome activation by measuring IL-1α, IL-1β, and IL-18 secretion (Fig. 3, S6). WT THP-1s treated with DMSO control released substantial amounts of IL-1α, IL-1β, and IL-18 when infected with WT Stm. In contrast, infected WT THP-1s treated with MCC950 secreted decreased levels of IL-1α, IL-1β, and IL-18, which are comparable to levels observed in WT Stm-infected *NAIP^-/-^* or *NLRC4^-/-^* THP-1s. (Fig. 3, S6A-D). Interestingly, WT Stm-infected *NAIP^-/-^* or *NLRC4^-/-^* THP-1s treated with MCC950 largely had significantly decreased IL-1α, IL-1β, and IL-18 secretion compared to infected *NAIP^-/-^* or *NLRC4^-/-^* THP-1s treated with DMSO or infected WT THP-1s treated with MCC950 (Fig. 3, S6A-D). Furthermore, *NAIP^-/-^* or *NLRC4^-/-^* THP-1s treated with MCC950 secreted negligible levels of IL-1α, IL-1β, and IL-18, similar to those observed during Δ*sipB* Stm infection (Fig. 3, S6A-D). WT, *NAIP^-/-^,* and *NLRC4^-/-^* THP-1s demonstrated robust IL-1α, IL-1β, and IL-18 secretion in response to LPS + nigericin that was significantly reduced by MCC950 treatment, indicating that this inhibitor effectively blocked NLRP3 inflammasome activation, as expected (Fig. 3, S6A-D). Release of the inflammasome-independent cytokine TNF-α was similar across the various THP-1 genotypes and treatments following infection (Fig. S6E, S6F). Altogether, these data indicate that *Salmonella* infection induces both NAIP/NLRC4- and NLRP3- dependent inflammasome activation in human macrophages.

**Fig 3.**
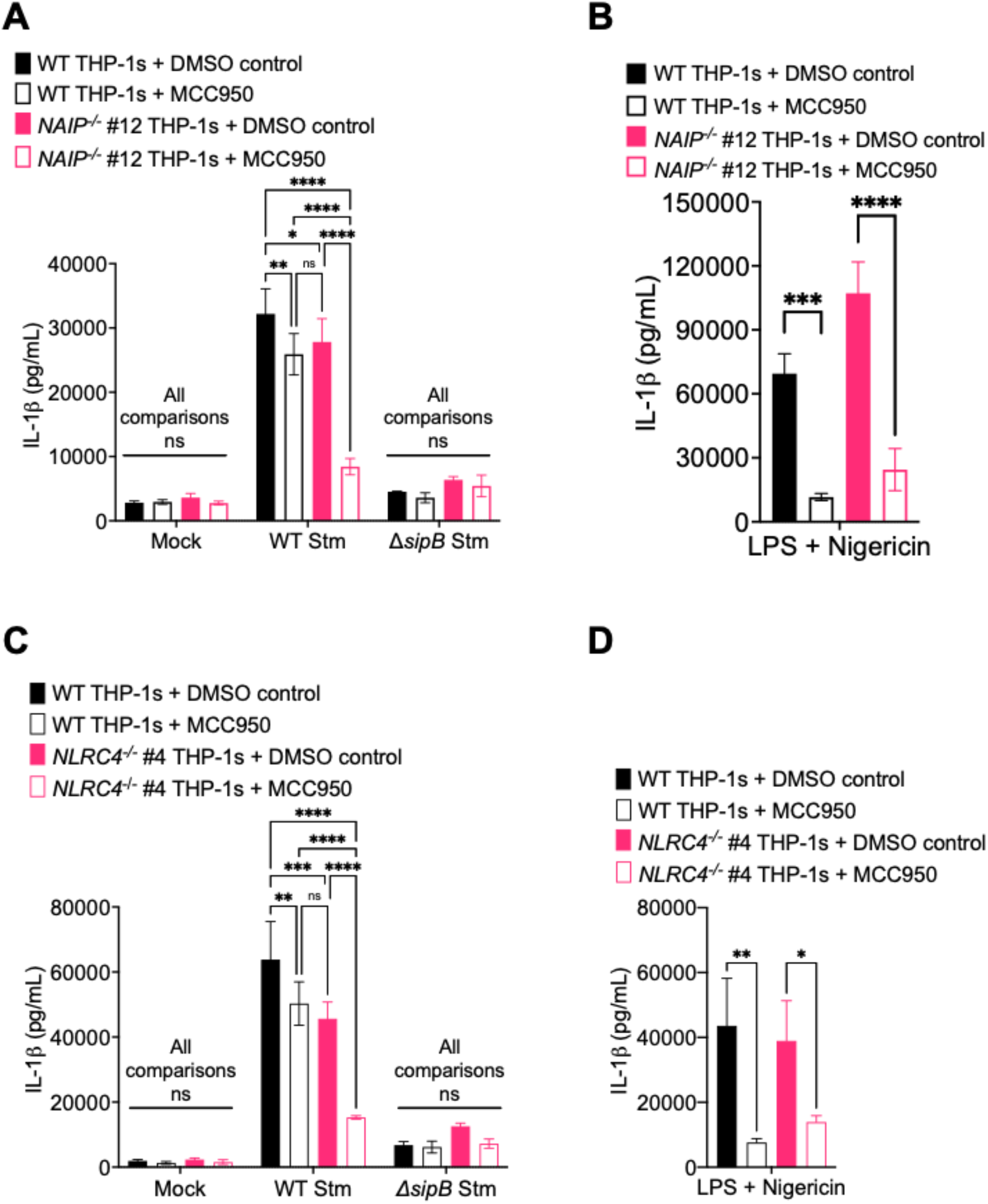
*Salmonella* induces NAIP/NLRC4- and NLRP3-dependent inflammasome activation in human macrophages. WT, *NAIP^-/-^*, or *NLRC4^-/-^* THP-1 monocyte-derived macrophages were primed with 100 ng/mL Pam3CSK4 for 16 hours. One hour prior to infection, cells were treated with 1 µM MCC950, a chemical inhibitor of the NLRP3 inflammasome or DMSO as a control. Cells were then infected with PBS (Mock), WT *S*. Typhimurium, or Δ*sipB S*. Typhimurium for 6 hours. As a control, cells were primed with 500 ng/mL LPS for 4 hours and treated with 10 µM nigericin for 6 hours. Release of IL- 1β into the supernatant was measured by ELISA. ns – not significant, \**p* < 0.05, \*\**p* < 0.01, \*\*\**p* < 0.001, \*\*\*\**p* < 0.0001 by Tukey’s multiple comparisons test (A, C) or by Šídák’s multiple comparisons test (B, D). Data shown are representative of at least three independent experiments.

### *Salmonella* induces NAIP/NLRC4- and CASP4/5-dependent inflammasome activation in human macrophages

In mice, in addition to the NAIP/NLRC4 and NLRP3 inflammasomes, *Salmonella* infection can also activate the caspase-11 inflammasome [41]. Caspase-11 detects cytosolic LPS and forms the noncanonical inflammasome, which secondarily activates the NLRP3 inflammasome [33, 42]. Caspases-4 and 5 are human orthologs of murine caspase-11 [33], and they can also sense cytosolic LPS to form the noncanonical inflammasome in human cells. We have previously observed caspase-4-dependent inflammasome activation in response to *Salmonella* infection in primary human macrophages [43], and caspases-4 and 5 also contribute to inflammasome responses to *Salmonella* infection in THP-1s and human intestinal epithelial cells [44, 45]. To test the relative contribution of both caspases-4 and 5 to NAIP-independent inflammasome responses during *Salmonella* infection of THP-1 macrophages, we treated WT or *NAIP^-/-^* THP-1s with siRNAs targeting *CASP4*, *CASP5,* or both, achieving ∼70-90% knockdown efficiency at the mRNA level (Fig. S7), and subsequently assayed for IL-1β secretion in response to WT Stm. WT THP-1s treated with either *CASP4* or *CASP5* siRNAs exhibited significantly decreased IL-1β secretion following WT Stm infection relative to WT THP-1s treated with control siRNA (Fig. 4A & B), in agreement with our previous observations in primary human macrophages [43]. *NAIP^-/-^* THP-1s treated with *CASP5* siRNA showed a slight but significant decrease in IL-1β secretion following *CASP5* siRNA treatment, but not *CASP4* siRNA treatment, compared to control siRNA-treated cells following WT Stm infection (Fig. 4A & B). WT and *NAIP^-/-^* THP-1s treated with both *CASP4* and *CASP5* siRNAs displayed significantly reduced IL-1β secretion relative to THP-1s treated with a scrambled control siRNA, although inflammasome activation was not completely abrogated when both *CASP4* and *CASP5* were knocked down in *NAIP^-/-^* THP-1 cells (Fig. 4C). As a control, we assessed inflammasome activation in response to transfected *E. coli* LPS, which activates the caspase-4/5 inflammasome. Both WT and *NAIP^-/-^* cells transfected with LPS displayed significantly decreased IL-1β secretion when *CASP4* was silenced, either alone or in conjunction with *CASP5* (Fig. 4A & C), whereas knockdown of *CASP5* alone did not significantly affect IL-1β secretion, as expected [45] (Fig. 4B). Taken together, these data suggest that the caspase-4/5 inflammasome is involved in the NAIP-independent response to *Salmonella*.

**Fig 4.**
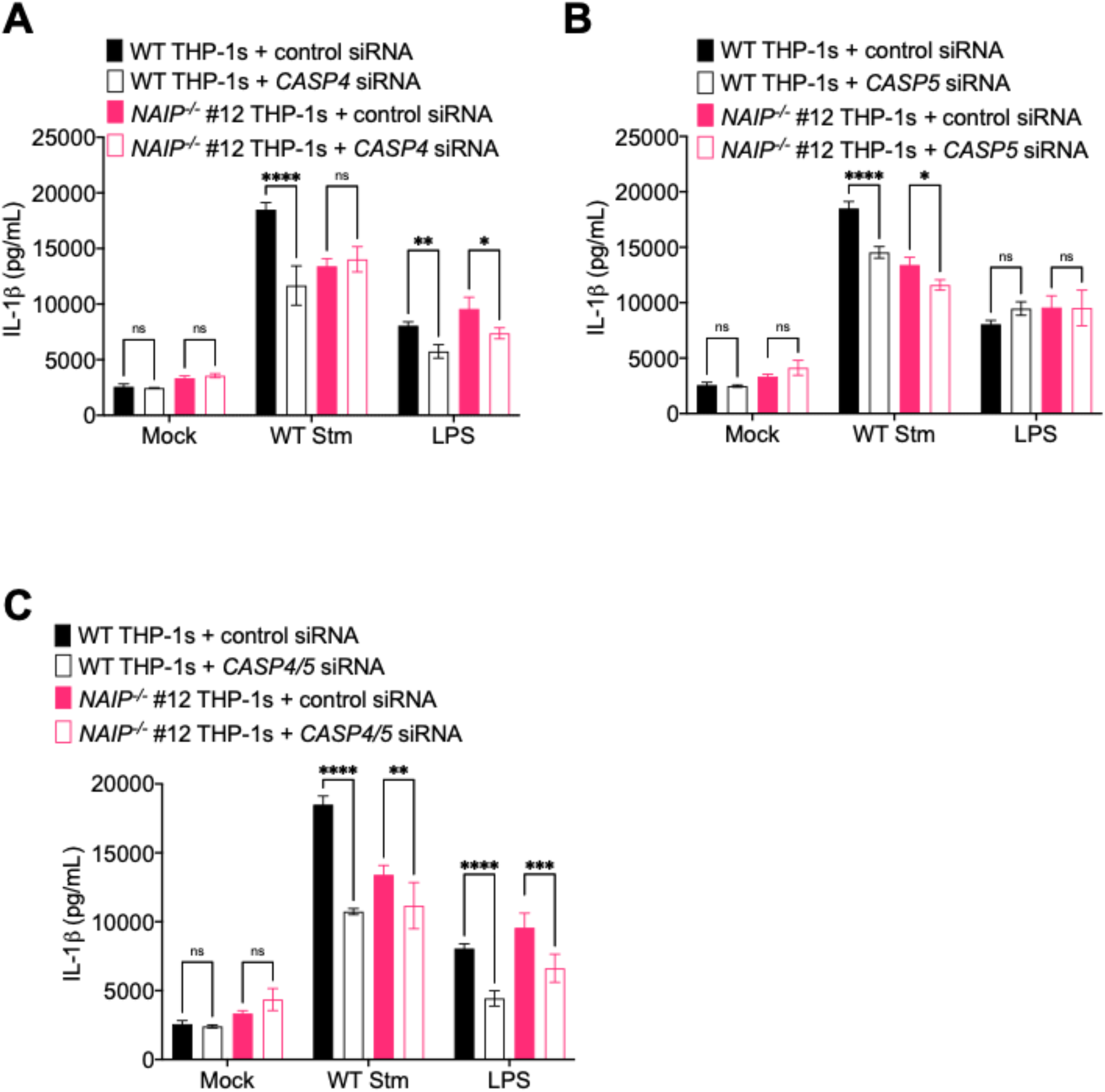
*Salmonella* induces NAIP- and CASP4/5-dependent inflammasome activation in human macrophages. WT or *NAIP^-/-^* THP-1 monocyte-derived macrophages were treated with siRNA targeting a control scrambled siRNA, siRNA targeting *CASP4* or *CASP5*, or siRNA targeting both *CASP4* and *CASP5* for 48 hours. Cells were primed with 100 ng/mL Pam3CSK4 for 16 hours. Cells were then infected with PBS (Mock) or WT *S*. Typhimurium for 6 hours. Release of IL-1β into the supernatant were measured by ELISA. As a control, cells were transfected with LPS. ns – not significant, \**p* < 0.05, \*\**p* < 0.01, \*\*\**p* < 0.001, \*\*\*\**p* < 0.0001 by Tukey’s multiple comparisons test. Data shown are representative of at least three independent experiments.

### The NAIP/NLRC4 and NLRP3 inflammasomes restrict *Salmonella* replication within human macrophages

One of the mechanisms by which inflammasome activation leads to control of bacterial infection is by restricting intracellular bacterial replication. In mice, the NAIP/NLRC4 inflammasome is important for controlling *Salmonella* replication in the intestine [28], whereas the NLRP3 inflammasome is dispensable for control of *Salmonella* infection *in vivo* [46, 47]. Caspases-1 and 11 restrict cytosolic *Salmonella* replication within murine macrophages [48]. Whether inflammasome activation restricts WT *Salmonella* replication in human macrophages is unknown. To test the hypothesis that inflammasome activation restricts *Salmonella* replication within human macrophages, we infected WT or *NAIP^-/-^* THP-1 macrophages with WT Stm in the presence or absence of the NLRP3 inhibitor MCC950 and determined the bacterial colony forming units (CFU) at various timepoints post-infection to assay bacterial replication. At 2 hours post-infection, we did not observe any differences in bacterial uptake between the different conditions (Fig. S8A). At 6 or 24 hours post-infection, the bacterial burden was the lowest in WT THP-1s, whereas *NAIP^-/-^* THP-1s harbored significantly higher bacterial burdens (Fig. 5A, S8B). WT THP-1s treated with MCC950 also contained a significantly higher number of bacterial CFUs, comparable to those in *NAIP^-/-^* THP-1s (Fig. 5A, S8B). *NAIP^-/-^* THP-1s treated with MCC950 had the highest bacterial burdens, which were significantly higher than the bacterial burdens in DMSO control-treated *NAIP^-/-^* THP-1s or WT THP-s treated with MCC950 (Fig. 5A). We then examined the fold-change in bacterial replication at 6 and 24 hours relative to 2 hours post-infection. The fold-change in bacterial replication was restricted the most effectively in WT THP-1s, moderately restricted in *NAIP^-/-^* THP-1s or WT THP-1s treated with MCC950, and the least restricted in *NAIP^-/-^* THP-1s treated with MCC950 (Fig. 5B). Collectively, these data suggest that both the NAIP/NLRC4 and NLRP3 inflammasomes restrict intracellular *Salmonella* replication within human macrophages at both early (6 hours post-infection) and late (24 hours post-infection) timepoints.

**Fig 5.**
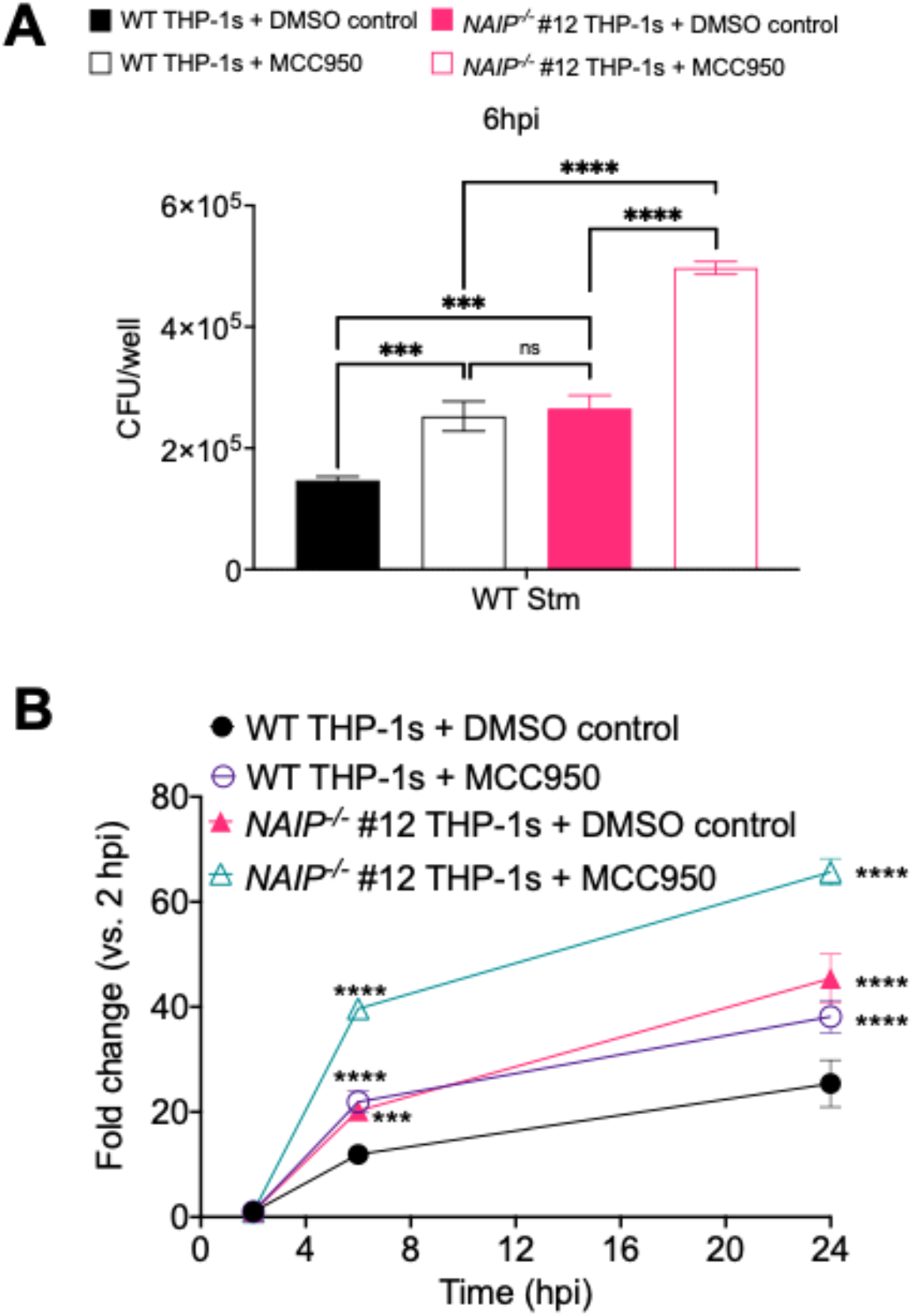
The NAIP and NLRP3 inflammasomes restrict *Salmonella* replication within human macrophages. WT or *NAIP^-/-^* THP-1 monocyte-derived macrophages were primed with 100 ng/mL Pam3CSK4 for 16 hours. One hour prior to infection, cells were treated with 1 µM MCC950, a chemical inhibitor of the NLRP3 inflammasome or DMSO as a control. Cells were then infected with PBS (Mock) or WT *S*. Typhimurium. Cells were lysed at the indicated time points and bacteria were plated to calculate CFU. (A) CFU/well of bacteria at 6 hpi (B) Fold change in CFU/well of bacteria at indicated time point, relative to 2 hpi CFU/well. ns – not significant, \*\*\**p* < 0.001, \*\*\*\**p* < 0.0001 by Dunnett’s multiple comparisons test (A) or Tukey’s multiple comparisons test (B). Data shown are representative of at least three independent experiments.

### *Salmonella* SPI-2 needle protein SsaG activates the NAIP/NLRC4 inflammasome in human macrophages

The *Salmonella* flagellin, SPI-1 T3SS inner rod (PrgJ), and needle (PrgI) proteins all activate NAIP in primary human macrophages, whereas the *Salmonella* SPI-2 T3SS inner rod protein (SsaI) is not sensed by NAIP [31, 32]. Similarly in mice, SsaI is not sensed by NAIP2 [11]. These findings have led to models proposing that the SPI-2 T3SS evades inflammasome detection to allow *Salmonella* to replicate or persist in both murine and human cells [11, 32]. However, our data indicate that the NAIP/NLRC4 inflammasome restricts *Salmonella* replication within macrophages even at late timepoints, when the SPI-1 T3SS and flagellin are thought to be downregulated [49–51]. As *Salmonella* utilizes the SPI-2 T3SS to replicate within macrophages [52], we asked whether the human NAIP/NLRC4 inflammasome detects the SPI-2 T3SS needle SsaG. To address this question, we delivered bacterial ligands into the cytosol of primary human monocyte-derived macrophages (hMDMs) derived from anonymous healthy human donors using the Gram-positive bacterium *Listeria monocytogenes*, which, upon infection, escapes from its vacuole into the cytosol where it expresses the protein ActA on its surface. Fusing bacterial ligands of interest to the N-terminus of truncated ActA allows these ligands to be delivered into the host cytosol, where they trigger NAIP/NLRC4 inflammasome activation [32, 53]. We infected hMDMs with WT *Listeria* (Lm) or *Listeria* expressing PrgJ, SsaI, or SsaG and assayed for inflammasome activation (Fig. 6A, S9). hMDMs infected with *Listeria* expressing the SPI-1 T3SS inner rod protein PrgJ induced robust inflammasome activation, indicated by significantly increased IL-18 secretion as well as robust IL-1α and IL-1β secretion compared to mock infection or WT Lm infection alone (Fig. 6A, S9), in agreement with our previous findings [32]. In contrast, and as we previously observed [32], *Listeria* expressing the SPI-2 inner rod protein SsaI failed to induce IL-1β, IL-18, and IL-1α secretion or cell death in hMDMs (Fig. 6A, S9). Intriguingly, we observed that *Listeria* expressing the SPI-2 needle protein SsaG induced significantly increased IL-18 and robust IL-1α and IL-1β secretion compared to mock infection or WT Lm infection alone (Fig. 6A, S9).

**Fig 6.**
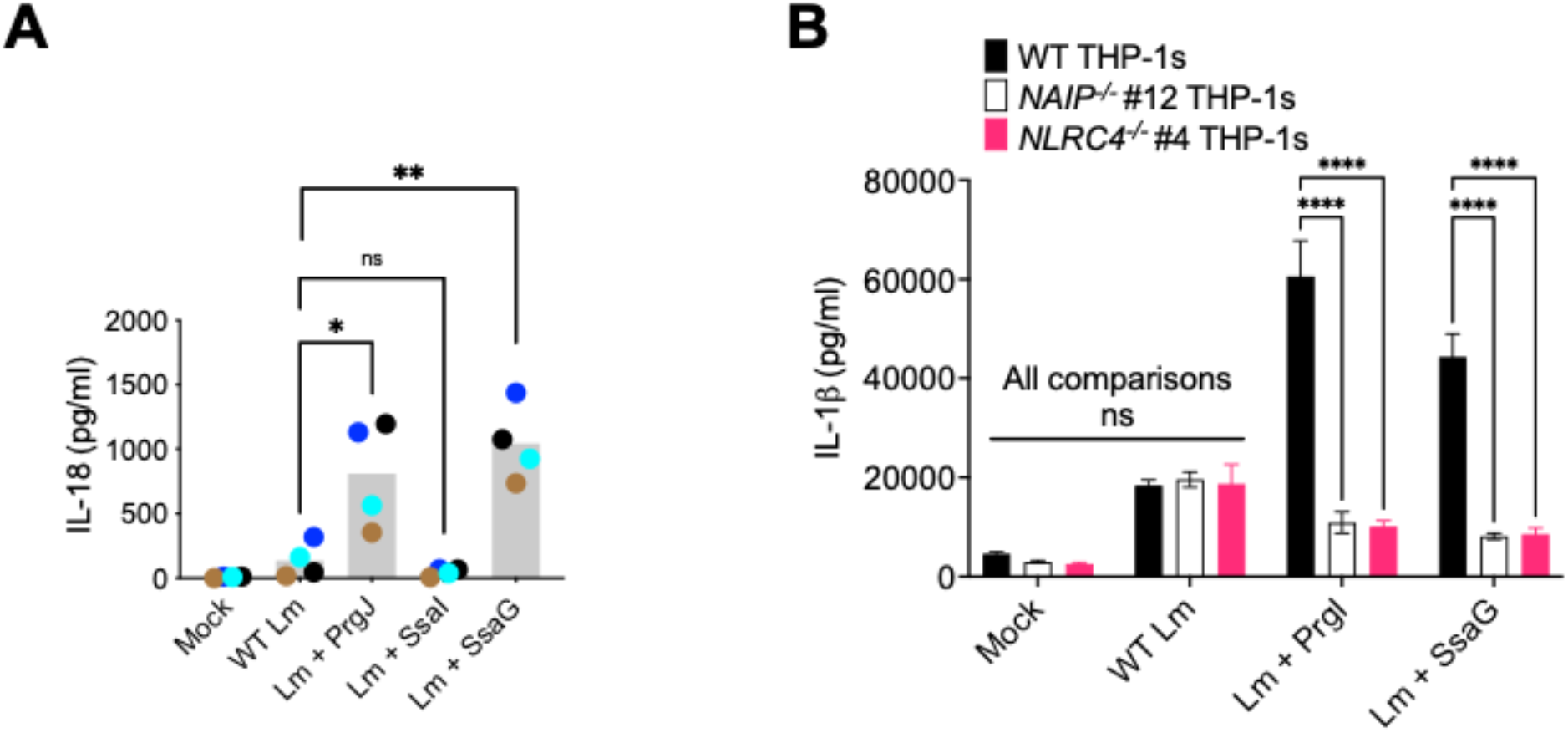
*Salmonella* SPI-2 needle protein SsaG activates the NAIP/NLRC4 inflammasome in human macrophages. (A) Primary hMDMs from four healthy human donors were infected with PBS (Mock), WT *Listeria (*WT Lm), *Listeria* expressing PrgJ (Lm + PrgJ), SsaI (Lm + SsaI), or SsaG (Lm + SsaG) for 16 hours at MOI=5. Release of IL-18 into the supernatant was measured by ELISA. Each dot represents the mean of individual donors derived from triplicate wells. The grey bar represents the mean of all donors. (B) WT or *NAIP^-/-^, NLRC4^-/-^* THP-1 monocyte-derived macrophages were primed with 100 ng/mL Pam3CSK4 for 16 hours. Cells were treated with PBS (Mock), WT *Listeria (*WT Lm), *Listeria* expressing PrgI (Lm + PrgI), or SsaG (Lm + SsaG) for 6 hours at MOI=20. Release of IL-1β into the supernatant was measured by ELISA. ns – not significant, * *p* < 0.05, ** *p* < 0.01, \*\*\*\**p* < 0.0001 paired t-test (A) or by Dunnett’s multiple comparisons test (B). Data shown are representative of at least three independent experiments.

To test whether NAIP or NLRC4 are required for inflammasome responses to SsaG, we infected WT, *NAIP^-/-^*, and *NLRC4^-/-^* THP-1s with WT *Listeria* (Lm) or *Listeria* expressing PrgI or SsaG and assayed for subsequent inflammasome activation by measuring levels of IL-1β, IL-18, and IL-1α secretion and cell death (Fig. 6B, S10). Infection of WT THP-1s with *Listeria* expressing PrgI or SsaG led to robust release of IL-1 cytokines and cytotoxicity. In contrast, *NAIP^-/-^* and *NLRC4^-/-^* THP-1s infected with *Listeria* expressing PrgI or SsaG released significantly reduced levels of IL-1 cytokines and cell death relative to WT THP-1s that were comparable to the background levels secreted by THP-1s infected with WT Lm (Fig. 6B, S10). Altogether, these data demonstrate that the SPI-2 needle protein activates the human NAIP/NLRC4 inflammasome, providing evidence that human NAIP can sense and respond to the *Salmonella* SPI-2 T3SS.

### NAIP/NLRC4 inflammasome recognition of the SPI-2 T3SS restricts *Salmonella* replication in human macrophages

To determine if NAIP/NLRC4-mediated recognition of the SPI-2 T3SS needle restricts *Salmonella* intracellular replication, we generated a *Salmonella* mutant strain (Δ*prgIfliCfljB*) lacking flagellin and the SPI-1 T3SS needle protein, PrgI. This strain is therefore unable to assemble a functional SPI-1 T3SS, but still expresses a functional SPI-2 T3SS. We infected WT or *NAIP^-/-^* THP-1 macrophages with Δ*prgIfliCfljB* and determined the CFUs at various timepoints to assay bacterial replication (Fig. 7, S11). Bacterial replication of Δ*prgIfliCfljB* over a 24-hour post-infection time course was restricted the most effectively in WT THP-1s and was significantly less restricted in *NAIP^-/-^* THP-1s (Fig. 7, S11). Collectively, our data suggest that there is SPI-1 T3SS/flagellin-independent, NAIP/NLRC4 inflammasome-dependent control of *Salmonella* replication in human macrophages, and that NAIP/NLRC4 recognition of the SPI-2 T3SS needle SsaG mediates such restriction of *Salmonella* replication in human macrophages.

**Fig 7.**
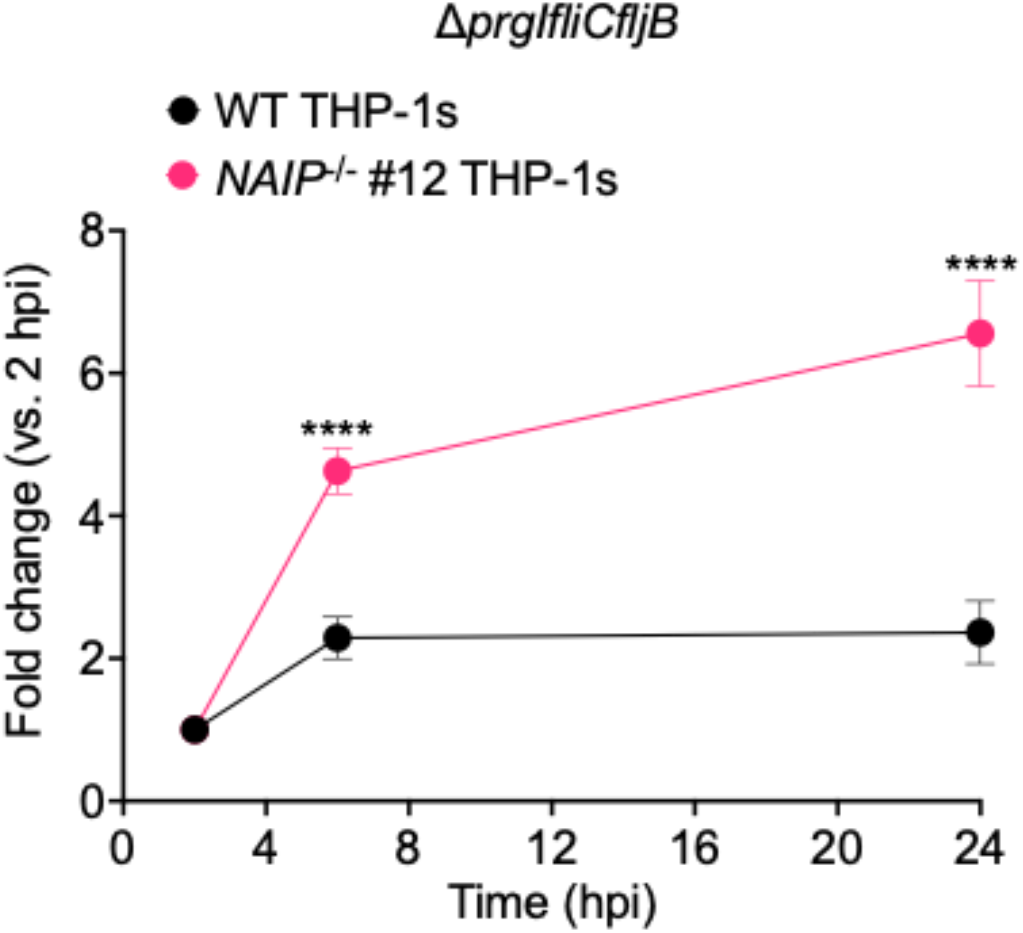
NAIP/NLRC4 inflammasome recognition of the SPI-2 T3SS restricts *Salmonella* replication in human macrophages. WT or *NAIP^-/-^* THP-1 monocyte-derived macrophages were primed with 100 ng/mL Pam3CSK4 for 16 hours. Cells were then infected with PBS (Mock) or Δ*prgIfliCfljB S*. Typhimurium. Cells were lysed at the indicated time points and bacteria were plated to calculate CFU. Fold change in CFU/well of bacteria at indicated time point, relative to 2 hpi CFU/well. ns – not significant, \*\*\**p* < 0.001, \*\*\*\**p* < 0.0001 by Tukey’s multiple comparisons test.

## Discussion

Our data show that human macrophages engage multiple inflammasome pathways to sense and respond to *Salmonella* infection. Using *NAIP^-/-^* and *NLRC4^-/-^* THP-1s (Fig. S1, S2), we found inflammasome activation in response to individual SPI-1 T3SS ligands to be entirely dependent on the NAIP/NLRC4 inflammasome in human macrophages (Fig. 1, S3, S4). In contrast, *Salmonella* infection induced activation of inflammasome responses that depended partially on NAIP/NLRC4, NLRP3, and CASP4/5 (Fig. 2-4, S5-7). Our findings are in agreement with a recent study demonstrating that both NLRC4 and NLRP3 are required for inflammasome responses to *Salmonella* in human macrophages [36]. Importantly, our data also reveal that both the NAIP/NLRC4 and NLRP3 inflammasomes contribute to restriction of *Salmonella* replication in human macrophages (Fig. 5, S8). Furthermore, contrary to the prevailing model that the SPI-2 T3SS evades NAIP detection, we show that the NAIP/NLRC4 inflammasome can recognize the *Salmonella* SPI-2 T3SS needle SsaG (Fig. 6, S9, S10), and that NAIP/NLRC4-dependent recognition of the SPI-2 T3SS restricts bacterial replication within human macrophages (Fig. 6, S11).

Many Gram-negative bacteria use evolutionarily conserved T3SSs to deliver virulence factors, or effectors, into host cells. We have previously shown that T3SS inner rod proteins from various Gram-negative bacteria activate the inflammasome in human macrophages [32]. In this study, we used T3SS inner rod, needle, or flagellin proteins from three different Gram-negative bacteria, *Salmonella*, *Burkholderia*, and *Legionella*, and observed that inflammasome activation in response to an isolated ligand is entirely dependent on NAIP/NLRC4 (Fig. 1, S3, S4, 6B). How the single human NAIP senses and responds to these diverse bacterial structures remains an open question. The *Salmonella* SPI-1 T3SS inner rod (PrgJ), SPI-1 T3SS needle (PrgI), and flagellin proteins exhibit low total sequence conservation, but they all retain several conserved hydrophobic amino acid residues within their structurally homologous C-terminal helices [11, 54]. In particular, both PrgJ and flagellin contain C-terminal leucine residues, specifically in their LLR motifs, which are critical for recognition by mNAIP2 and mNAIP5, respectively [55–59]. Instead of the LLR motif, PrgI has other hydrophobic amino acids, including valine and isoleucine residues, within its C-terminal helical domain. These terminal hydrophobic residues within PrgI are important for mediating inflammasome activation in human macrophages [17]. Interestingly, an alignment of the amino acid sequences of the SPI-2 T3SS needle protein (SsaG), PrgJ, and PrgI using Clustal Omega revealed that SsaG also contains conserved hydrophobic amino acid residues in its C-terminus (Fig S12A)). Specifically, SsaG has C-terminal isoleucine residues like PrgI. To further compare these ligands at the structural level, we examined published three-dimensional structures of PrgJ and PrgI and used PHYRE2 Protein Fold Recognition Server to predict the structure of SsaG. Similar to PrgJ and PrgI, SsaG also displays an alpha-helical structure at its C-terminus (Fig S12B)). Thus, SsaG displays secondary structural and sequence motifs similar to those retained by the other T3SS ligands recognized by human NAIP. Unlike these T3SS ligands, the *Salmonella* SPI-2 inner rod protein, SsaI, does not retain such conserved C-terminal residues. Perhaps this is why SsaI is not detected by human NAIP. Still, the specific ligand residues recognized by human NAIP remain unknown. Murine NAIPs use their nucleotide-binding domain (NBD)-associated domains to detect these conserved residues of their respective cognate bacterial ligand [55–60]. It remains to be determined if the single human NAIP uses a similar mechanism to broadly detect its bacterial ligands.

Human NAIP is a generalist, as it detects multiple bacterial ligands, while the murine NAIPs are specialists, as they each recognize a particular ligand. The functional consequences of being a generalist NAIP is unclear. It is possible that recognizing a broad array of structures diminishes the affinity with which human NAIP binds its ligands. Alternatively, human NAIP may recognize its bacterial ligands with varying affinities. Furthermore, under physiological conditions, all bacterial ligands may not be delivered to the cytosol to the same extent or recognized with the same sensitivity. Varying levels of inflammasome activation with the different ligands may have distinct downstream consequences. It would be interesting to determine whether restriction of bacterial replication varies depending on which bacterial ligand is sensed.

*Salmonella* infection induces NAIP/NLRC4-, CASP4/5-, and NLRP3-dependent inflammasome activation in human macrophages (Fig. 2-4, S5-7). This suggests that there is redundancy in the inflammasome pathways when sensing and responding to *Salmonella* infection, such that loss of just one inflammasome does not result in severe loss of inflammasome activation in human macrophages. Given our observations with individual ligand delivery (Fig. 1, S3, S4), it is likely that the NAIP/NLRC4 inflammasome is sensing the *Salmonella* SPI-1 T3SS inner rod, SPI-1 and SPI-2 needle, and flagellin proteins during infection. However, it remains unknown how NLRP3 and CASP4/5 inflammasomes are activated in human macrophages during *Salmonella* infection. CASP4/5 detects intracellular LPS [33], but given that *Salmonella* is normally a vacuolar pathogen in macrophages, it is unclear how CASP4/5 may be accessing LPS. In murine macrophages, a small percentage of *Salmonella-*containing vacuoles rupture, allowing bacteria to escape into the host cell cytosol [61]. In human intestinal epithelial cells, a subpopulation of *Salmonella* that escape the vacuole and replicates in the cytosol activates the CASP4/5 inflammasome at late timepoints of infection [44]. Moreover, other host immune factors can potentiate inflammasome signaling by promoting the release of PAMPs, such as LPS, into the host cell cytosol. For example, a family of host immune factors called guanylate binding proteins (GBPs) can localize to pathogen-containing vacuoles [62]. Murine GBPs promote rupture of the *Salmonella*-containing vacuole (SCV) [61]. Human GBP-1 can localize to the SCV in macrophages [63], and in human epithelial cells, GBP1 binds to bacterial LPS on the surface of cytosolic *Salmonella* and promotes the recruitment and activation of caspase-4 [64, 65]. Another mechanism by which LPS can access the cytosol is through bacterial outer membrane vesicles (OMVs), and this mechanism has been shown to activate the caspase-11 inflammasome in murine models [66, 67]. Future studies will explore if the CASP4/5- dependent inflammasome activation we have observed in human macrophages is facilitated by *Salmonella* escape into the host cell cytosol, GBP1 activity, OMVs, or other mechanisms.

The NLRP3 inflammasome can be activated by a variety of different stimuli, including potassium efflux. It can also be activated downstream of the CASP4/5 inflammasome [42, 45], leading to non-canonical NLRP3 inflammasome activation. Given that we observed only partial loss of inflammasome activation in the *NAIP^-/-^* THP- 1s treated with siRNA targeting *CASP4* and *CASP5*, we hypothesize that at least part of the NLRP3-dependent response is due to canonical activation (Fig. 4), although this partial loss may also be due to incomplete knockdown of *CASP4* and *CASP5*. A recent study also found that *Salmonella* infection induces NLRC4- and NLRP3-dependent inflammasome activation in human macrophages, and observed that full-length *Salmonella* flagellin can activate the NLRP3 inflammasome [36]. In contrast, we found the response to flagellin to be entirely dependent on the NAIP/NLRC4 inflammasome (Fig. 1, S3, S4). The reason for this apparent discrepancy is unclear, but in our studies, we used a truncated flagellin that only contains the C-terminal D0 domain and thus does not stimulate TLR5 signaling [56, 68]. It is possible that full-length flagellin, in addition to activating the NAIP/NLRC4 inflammasome, also stimulates TLR5 signaling, perhaps potentiating NLRP3-dependent responses.

We observed NAIP/NLRC4- and NLRP3-dependent restriction of *Salmonella* (Fig. 5, S8), but the mechanism by which inflammasome activation promotes bacterial restriction is unclear. Inflammasome activation often triggers host cell death, thereby eliminating the pathogen’s intracellular replicative niche. *In vivo*, pyroptosis can trigger formation of pore-induced intracellular traps (PITs). These PITs can trap intracellular bacteria that can subsequently be efferocytosed by neutrophils [69]. However, in murine macrophages, inhibition of *Salmonella* replication by caspase-1 and caspase-11 occurs prior to host cell death, indicating that caspase-1 and caspase-11 restrict *Salmonella* through a mechanism distinct from cell death [48]. Another mechanism of inflammasome-dependent restriction may be through promoting phagolysomal maturation. In murine macrophages infected with *Legionella*, NAIP5 activation results in increased colocalization of *Legionella-*containing vacuoles with the lysosomal markers cathepsin-D and Lamp-1 [70, 71]. Perhaps a similar process occurs during *Salmonella* infection of human macrophages.

The current model is that the SPI-2 T3SS subverts inflammasome activation to facilitate *Salmonella*’s intracellular survival, based on the observation that the SPI-2 T3SS inner rod SsaI is not detected in murine or human macrophages [11, 32]. Moreover, the SPI-2 T3SS effectors are critical for biogenesis and maintenance of the SCV [52]. Thus, evasion of inflammasome activation by the SPI-2 T3SS was thought to confer an advantage to the pathogen. However, our findings indicate that the NAIP/NLRC4 inflammasome detects the *Salmonella* SPI-2 T3SS needle protein SsaG. Furthermore, we find that SPI-1-independent, flagellin-independent, NAIP-dependent detection of *Salmonella* mediates restriction of intracellular bacterial replication in human macrophages. Perhaps the NAIP/NLRC4-mediated detection of SsaG is a consequence of a functional constraint placed upon SsaG’s role as a T3SS needle protein. It is thus possible that SsaG is unable to evade immune detection due to such functional constraints.

While we focused here primarily on inflammasome responses in human macrophages, *Salmonella*’s first cellular encounters are with intestinal epithelial cells. In mice, NAIP/NLRC4 inflammasome activation in intestinal epithelial cells results in extrusion of infected cells from the epithelial layer [27, 28]. It has been proposed that this mechanism eliminates *Salmonella* from the host and helps control bacterial burdens. Whether similar NAIP/NLRC4-dependent mechanisms are engaged in human intestinal epithelial cells remains to be elucidated.

Overall, these data indicate that *Salmonella* infection of human macrophages triggers activation of multiple inflammasomes, and at least two of these inflammasomes, the NAIP/NLRC4, and the NLRP3 inflammasomes, appear to be essential for controlling bacterial replication within macrophages. Furthermore, our data indicate that the human NAIP/NLRC4 inflammasome detects the SPI-2 needle protein SsaG, and that NAIP/NLRC4-mediated detection of the SPI-2 T3SS restricts *Salmonella* replication within macrophages. Collectively, our findings provide fundamental insight into how *Salmonella* is sensed and restricted by human macrophages. Moreover, these results offer a foundation for further understanding of how each of these pathways is activated and how these inflammasomes interact to mediate downstream responses that promote control of *Salmonella* infection in human macrophages.

## Materials and Methods

### Ethics statement

All studies involving primary human monocyte-derived macrophages (hMDMs) were performed in compliance with the requirements of the US Department of Health and Human Services and the principles expressed in the Declaration of Helsinki. hMDMs were derived from samples obtained from the University of Pennsylvania Human Immunology Core. These samples are considered to be a secondary use of deidentified human specimens and are exempt via Title 55 Part 46, Subpart A of 46.101 (b) of the Code of Federal Regulations.

### Bacterial strains and growth conditions

Targeted deletion strains used in this study were made on the *Salmonella enterica* serovar Typhimurium SL1344 strain background. The Δ*prgIfliCfljB* strain was engineered using the Δ*fliCfljB* background [72], in which the SPI-1 T3SS needle, *prgI*, was deleted through a chloramphenicol resistance cassette insertion into *prgI* (*fliCfljBprgI*::CmR) using standard methods [73].

WT, Δ*sipB* [74], and Δ*prgIfliCfljB* isogenic strains were routinely grown overnight in Luria-Bertani (LB) broth with streptomycin (100 μg/ml) at 37°C. For infection of cultured cells, overnight cultures were diluted in LB containing 300 mM NaCl and grown standing for 3 hours at 37°C to induce SPI-1 expression [75].

*Listeria monocytogenes* WT and isogenic strains on the 10403S background were cultured in brain heart infusion (BHI) medium [53]. The *Listeria* strain encoding the heterologous bacterial ligand *S.* Typhimurium PrgJ translationally fused to the truncated N-terminus of ActA and under the control of the *actA* promoter was used [53]. The *Listeria* strains expressing *S.* Typhimurium SsaI and SsaG were constructed using codon-optimized gene fragments (IDT) cloned into the pPL2 vector and introduced into *Listeria* as previously described [53, 76].

### Cell culture of THP-1s

THP-1 cells (TIB-202; American Type Culture Collection) were maintained in RPMI supplemented with 10% (vol/vol) heat-inactivated FBS, 0.05 nM β- mercaptoethanol, 100 IU/mL penicillin, and 100 μg/mL streptomycin at 37°C in a humidified incubator. Two days before experimentation, the cells were replated in media without antibiotics in a 48-well plate at a concentration of 2 × 10^5^ cells/well and incubated with phorbol 12-myristate 13-acetate (PMA) for 24 hours to allow differentiation into macrophages. Macrophages were primed with 100 ng/mL Pam3CSK4 (Invivogen) for 16 hours prior to bacterial infections or anthrax toxin treatments. For experiments involving LPS, cells were pretreated with 500 ng/mL LPS (Sigma-Aldrich) for 3 hours. For experiments involving Nigericin, cells were treated with 10 μM Nigericin (EMD Millipore) for 6 hours. For experiments involving MCC950, cells were treated with 1 μM MCC950 (Sigma Aldrich) 1 hour prior to infection.

### Cell culture of primary human monocyte-derived macrophages (hMDMs)

Purified human monocytes from de-identified healthy human donors were obtained from the University of Pennsylvania Human Immunology Core. Monocytes were cultured in RPMI supplemented with 10% (vol/vol) heat-inactivated FBS, 2 mM L-glutamine, 100 IU/mL penicillin, 100 μg/ml streptomycin, and 50 ng/ml recombinant human M-CSF (Gemini Bio-Products) for 6 days to promote differentiation into hMDMs. One day prior to infection, adherent hMDMs were replated in media with 25 ng/ml human M-CSF lacking antibiotics at 1.0 × 10^5^ cells/well in a 48-well plate.

### Bacterial infections

Overnight cultures of *Salmonella* were diluted into LB broth containing 300 mM NaCl and grown for 3 hours standing at 37°C to induce SPI-1 expression [75]. Overnight cultures of *L. monocytogenes* were diluted and grown for 3 hours in BHI. All cultures were pelleted at 6,010 × *g* for 3 minutes, washed once with PBS, and resuspended in PBS. THP-1 cells were infected with *S.* Typhimurium or *L. monocytogenes* at a multiplicity of infection (MOI) of 20. hMDMs were infected with *L. monocytogenes* at an MOI of 5. Infected cells were centrifuged at 290 × *g* for 10 min and incubated at 37°C. 1 hour post-infection, cells were treated with 100 ng/mL or 50 ng/mL of gentamicin to kill any extracellular *S.* Typhimurium or *L. monocytogenes* respectively. *Salmonella* and *Listeria* infections in THP-1s proceeded at 37°C for 6 hours. *Listeria* infection of hMDMs proceeded at 37°C for 16 hours. For all experiments, control cells were mock-infected with PBS.

### Anthrax toxin-mediated delivery of bacterial ligands

Recombinant proteins (PA, LFn-FlaA^310-475^, LFn-PrgJ, and LFn-YscF) were kindly provided by Russell Vance [18]. PA and LFn doses for *in vitro* delivery were: 1 μg/ml PA for FlaTox; 4 μg/ml PA for PrgJTox and YscFTox; 500 ng/ml LFn-FlaA^310-475^; 8 ng/ml LFn-PrgJ; and 200 ng/mL LFn-YscF.

### siRNA-mediated knockdown of genes

All Silencer Select siRNA oligos were purchased from Ambion (Life Technologies). For *CASP4*, siRNA ID# s2412 was used. For *CASP5*, siRNA ID# s2417 was used. The two Silencer Select negative control siRNAs (Silencer Select Negative Control No. 1 siRNA and Silencer Select Negative Control No. 2 siRNA) were used as a control. Two days before infection, 30 nM of siRNA was transfected into macrophages using Lipofectamine RNAiMAX transfection reagent (Thermo Fisher Scientific) following the manufacturer’s protocol. 16 hours before infection, the media was replaced with fresh antibiotic-free media containing 100 ng/ml Pam3CSK4. In parallel, siRNA-transfected cells were also transfected with 2 μg/ml of *E. coli* LPS strain W3110 (kindly provided by Robert Ernst) using FuGENE HD transfection reagent (Promega) for 6 hours.

### Bacterial intracellular replication assay

Cells were infected with WT or Δ*prgIfliCfljB S.* Typhimurium as usual at an MOI of 20. 1 hour post-infection, cells were treated with 100 μg/ml of gentamicin to kill any extracellular bacteria. 2 hours post-infection, the media was replaced with fresh media containing 10 μg/ml of gentamicin. At the indicated time points, cells were lysed with PBS containing 0.5% Triton to collect all intracellular bacteria. Harvested bacteria were serially diluted in PBS and plated on LB agar with streptomycin (100 μg/ml) plates to enumerate colony forming units (CFUs). Plates were incubated at 37°C overnight and then CFUs were counted.

### ELISAs

Harvested supernatants from infected cells were assayed using ELISA kits for human IL-1α (R&D Systems), IL-18 (R&D Systems), IL-1β (BD Biosciences), and TNF-α (R&D Systems).

### LDH cytotoxicity assays

Harvested supernatants from infected cells were assayed for cytotoxicity by measuring loss of cellular membrane integrity via lactate dehydrogenase (LDH) activity. LDH release was quantified using an LDH Cytotoxicity Detection Kit (Clontech) according to the manufacturer’s instructions and normalized to mock-infected cells.

### Quantitative RT-PCR analysis

RNA was isolated using the RNeasy Plus Mini Kit (Qiagen) following the manufacturer’s instructions. Cells were lysed in 350 μL RLT buffer with β- mercaptoethanol and centrifuged through a QIAshredder spin column (Qiagen). cDNA was synthesized from isolated RNA using SuperScript II Reverse Transcriptase (Invitrogen) following the manufacturer’s protocol. Quantitative PCR was conducted with the CFX96 real-time system from Bio-Rad using the SsoFast EvaGreen Supermix with Low ROX (Bio-Rad). For analysis, mRNA levels of siRNA-treated cells were normalized to housekeeping gene *HPRT* and control siRNA-treated cells using the 2^−ΔΔCT^ (cycle threshold) method [77] to calculate knockdown efficiency. The following primers from PrimerBank were used. The PrimerBank identifications are *CASP4* (73622124c1), and *CASP5* (209870072c2), and *HPRT* (164518913c1); all 5′–3′:

*CASP4* forward: CAAGAGAAGCAACGTATGGCA

*CASP4* reverse: AGGCAGATGGTCAAACTCTGTA

*CASP5* forward: TTCAACACCACATAACGTGTCC

*CASP5* reverse: GTCAAGGTTGCTCGTTCTATGG

*HPRT* forward: CCTGGCGTCGTGATTAGTGAT

*HPRT* reverse: AGACGTTCAGTCCTGTCCATAA

### Statistical analysis

Prism 9.1.1 (GraphPad Software) was utilized for the graphing of data and all statistical analyses. Statistical significance for experiments with THP-1 cells was determined using the appropriate test and are indicated in each figure legend. Differences were considered statistically significant if the *p* value was <0.05.

## Acknowledgements

We thank members of Igor Brodsky’s and Sunny Shin’s laboratories for scientific discussion. We thank Meghan Wynosky-Dolfi for technical advice. We thank Russell Vance, Randilea Nichols, Isabella Rauch, and Jeannette Tenthorey for providing anthrax toxin-based reagents, JD Sauer for providing the *Listeria* strains and constructs for generating ActA fusion proteins, and Robert Ernst for providing *E. coli* LPS. We thank the Human Immunology Core of the Penn Center for AIDS Research and Abramson Cancer Center for providing purified primary human monocytes.

## Supporting Information

**S1 Fig.**
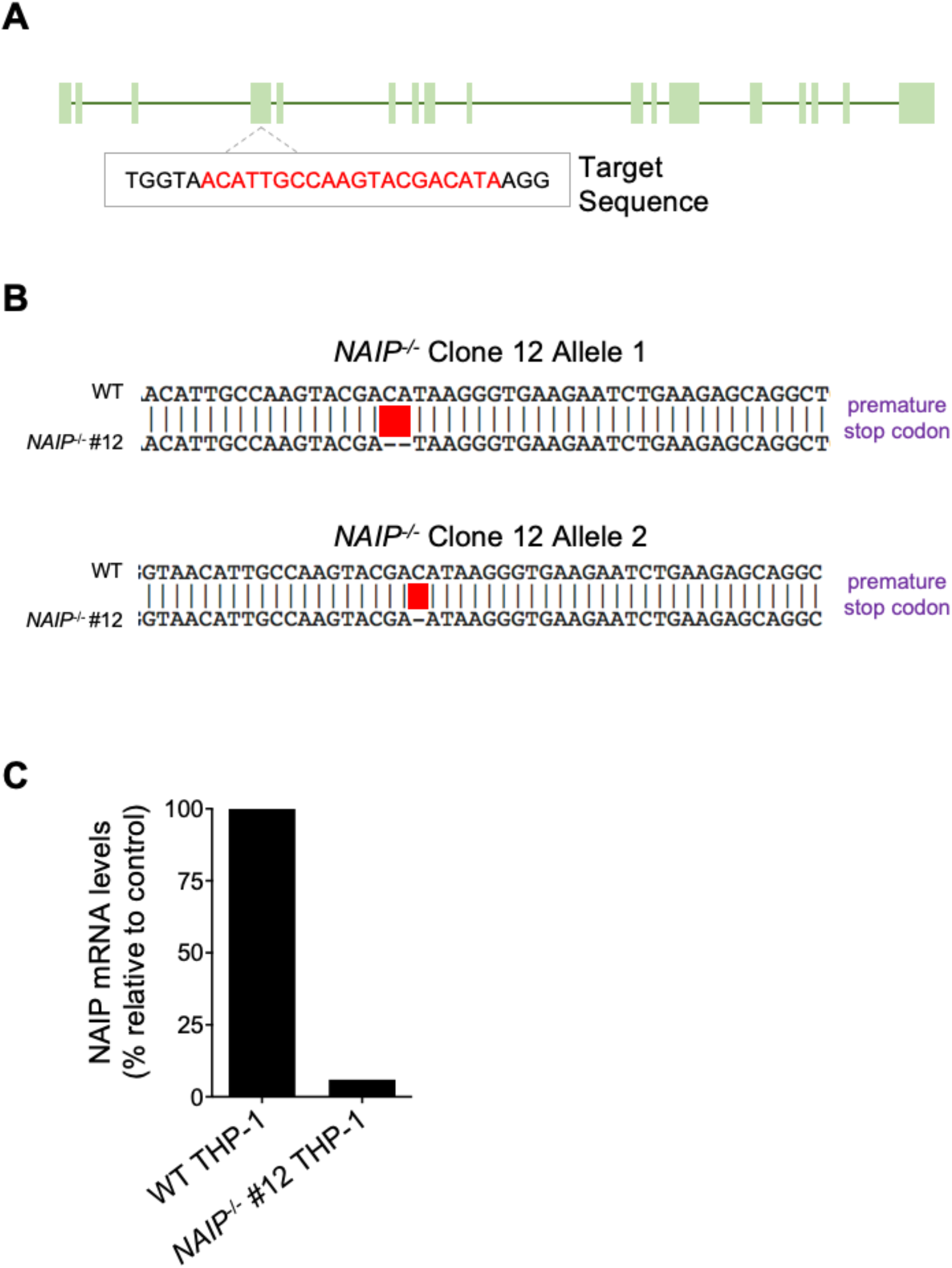
Validation of *NAIP* mutant THP-1 single cell clones generated with CRISPR/Cas9 genome editing. (A) Schematic representation of the *NAIP* gene with exons (filled boxes) and introns (filled lines). gRNA target sequence is highlighted in red. (B) Sequence alignments of WT THP-1 and *NAIP^-/-^* clone 12 are shown for both alleles. Red boxes represent the mutated region. Purple text represents the predicted impact of the mutation on the amino acid sequence. (C) qRT-PCR was performed to quantitate *NAIP* mRNA levels in WT THP-1 and *NAIP^-/-^* THP-1 cells. For the *NAIP^-/-^* THP-1 cells*, NAIP* mRNA levels were normalized to human HPRT mRNA levels and WT THP-1 cells.

**S2 Fig.**
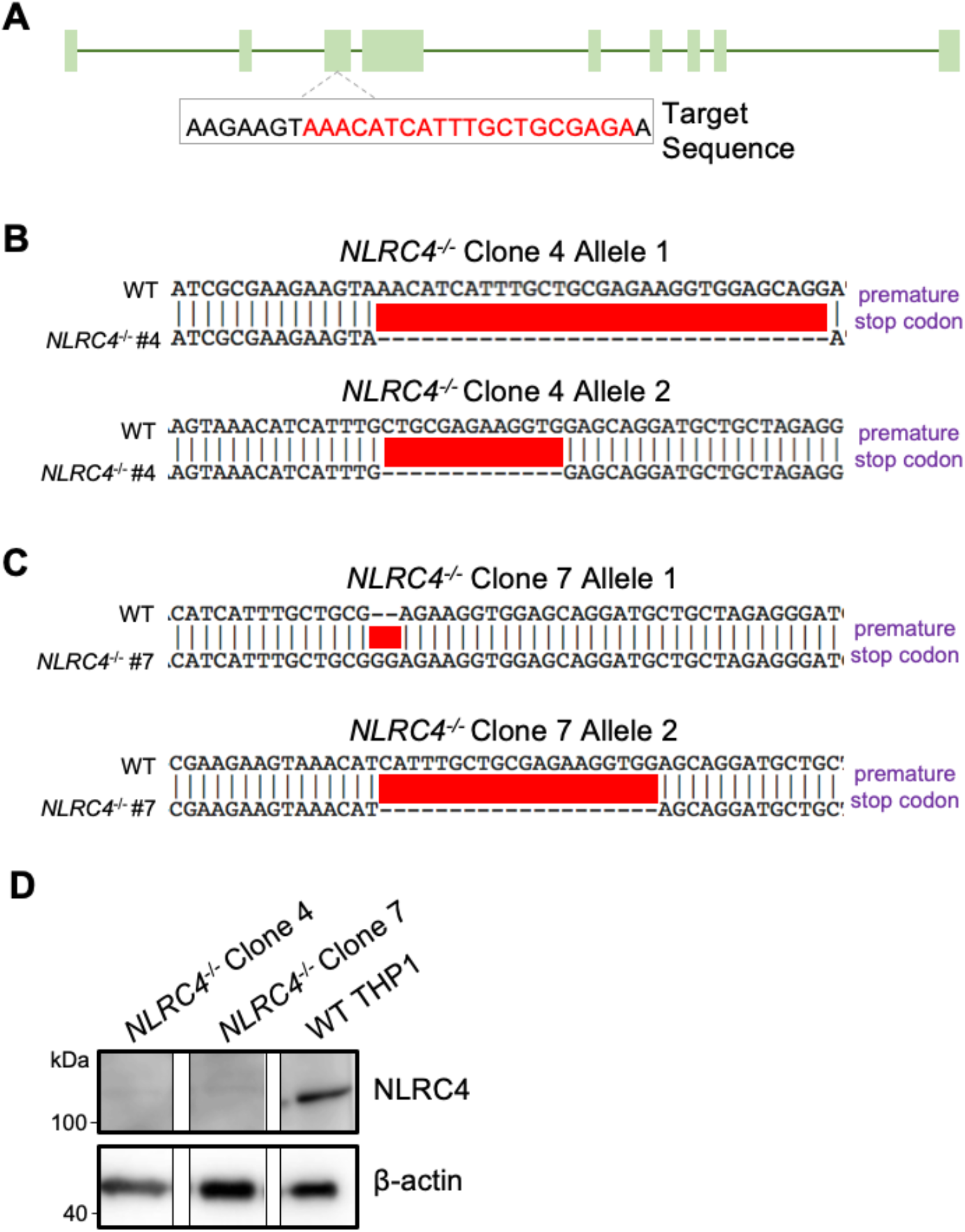
Validation of *NLRC4* mutant THP-1 single cell clones generated with CRISPR/Cas9-mediated genome editing. (A) Schematic representation of the *NLRC4* gene with exons (filled boxes) and introns (lines). gRNA target sequence is highlighted in red. (B-C) Sequence alignments of WT THP-1 and *NLRC4^-/-^* clones are shown for both alleles per clone. Red boxes highlight the mutated region. Purple text represents the predicted impact of the mutation on the amino acid sequence. (D) Immunoblot analysis was performed on cell lysates for human NLRC4, and β-actin as a loading control.

**S3 Fig. (related to Figure 1).**
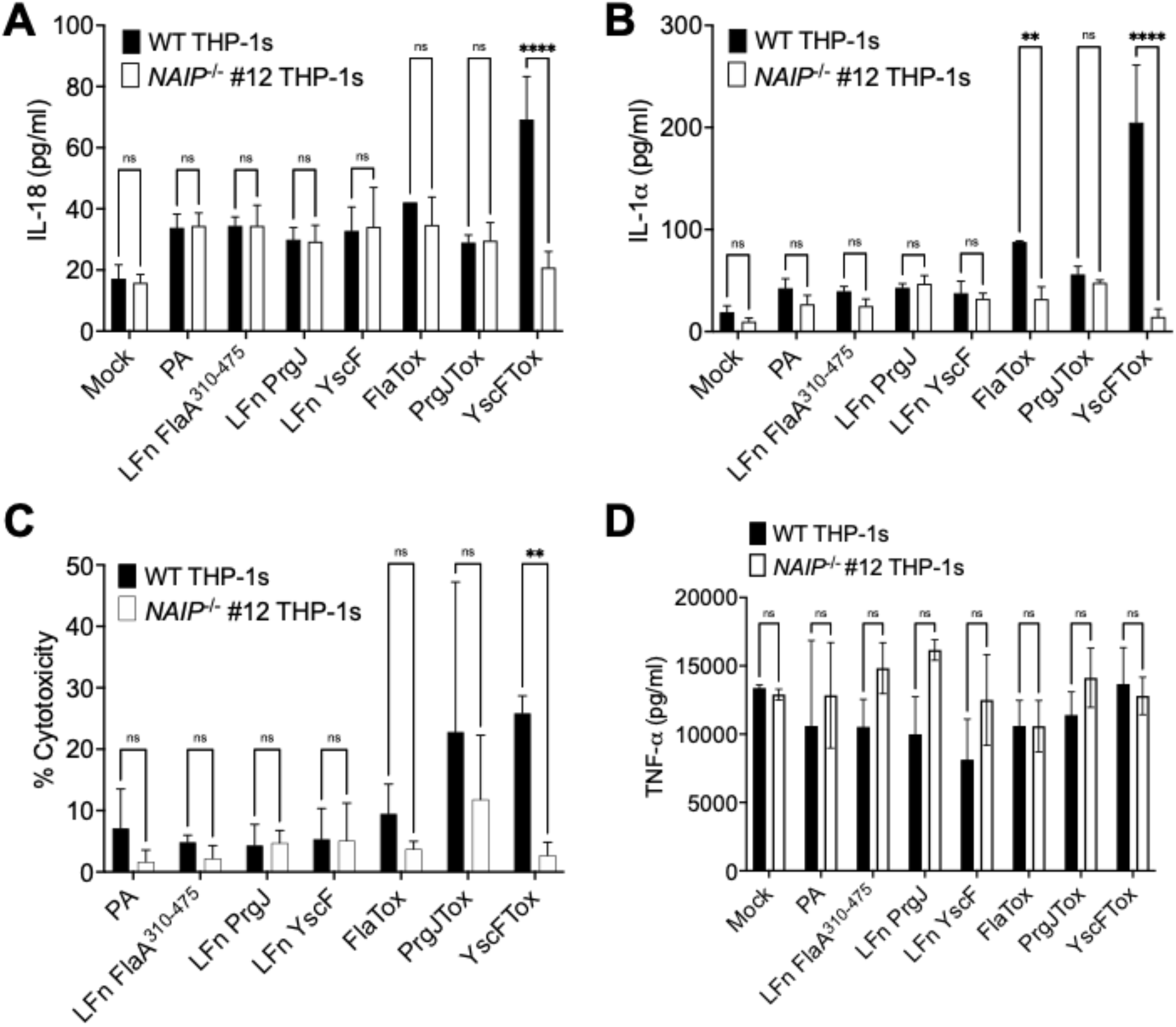
NAIP is necessary for inflammasome responses to T3SS ligands in human macrophages. WT or *NAIP^-/-^* THP-1 monocyte-derived macrophages were primed with 100 ng/ml Pam3CSK4 for 16 hours. Cells were then treated with PBS (Mock), PA alone, LFn FlaA^310–475^ alone, LFn PrgJ alone, LF YscF alone, PA+LFn FlaA^310–475^ (FlaTox), PA+LFn PrgJ (PrgJTox), or PA+LFn YscF (YscFTox) for 6 hours. (A, B, D) Release of cytokines IL-18, IL-1α, and TNF-α into the supernatant were measured by ELISA. (C) Cell death (percentage cytotoxicity) was measured by lactate dehydrogenase release assay and normalized to Mock-treated cells. ns – not significant, \*\**p* < 0.01, \*\*\*\**p* < 0.0001 by Šídák’s multiple comparisons test. Data shown are representative of at least three independent experiments.

**S4 Fig. (related to Figure 1).**
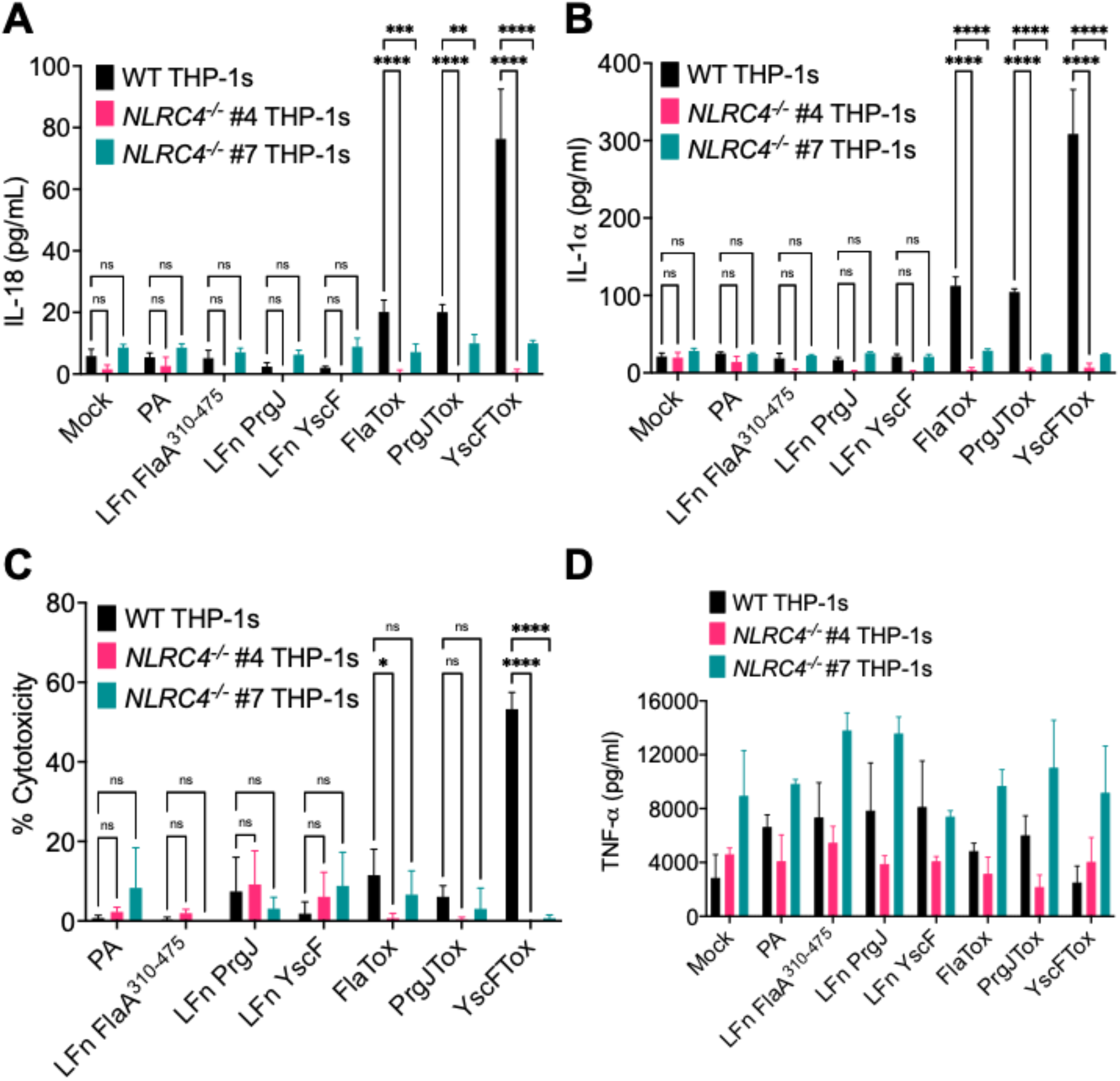
NLRC4 is necessary for inflammasome responses to T3SS ligands in human macrophages. WT or two independent clones of *NLRC4^-/-^* THP-1 monocyte-derived macrophages were primed with 100 ng/ml Pam3CSK4 for 16 hours. Cells were then treated with PBS (Mock), PA alone, LFn FlaA^310–475^ alone, LFn PrgJ alone, LFn YscF alone, PA+LFn FlaA^310–475^ (FlaTox), PA+LFn PrgJ (PrgJTox), or PA+LFn YscF (YscFTox) for 6 hours. (A, B, D) Release of cytokines IL-18, IL-1α, and TNF-α into the supernatant were measured by ELISA. (C) Cell death (percentage cytotoxicity) was measured by lactate dehydrogenase release assay and normalized to Mock-treated cells. ns – not significant, \**p* < 0.05, \*\**p* < 0.01, \*\*\**p* < 0.001, \*\*\*\**p* < 0.0001 by Dunnett’s multiple comparisons test (A-C). Data shown are representative of at least three independent experiments.

**S5 Fig. (related to Figure 2).**
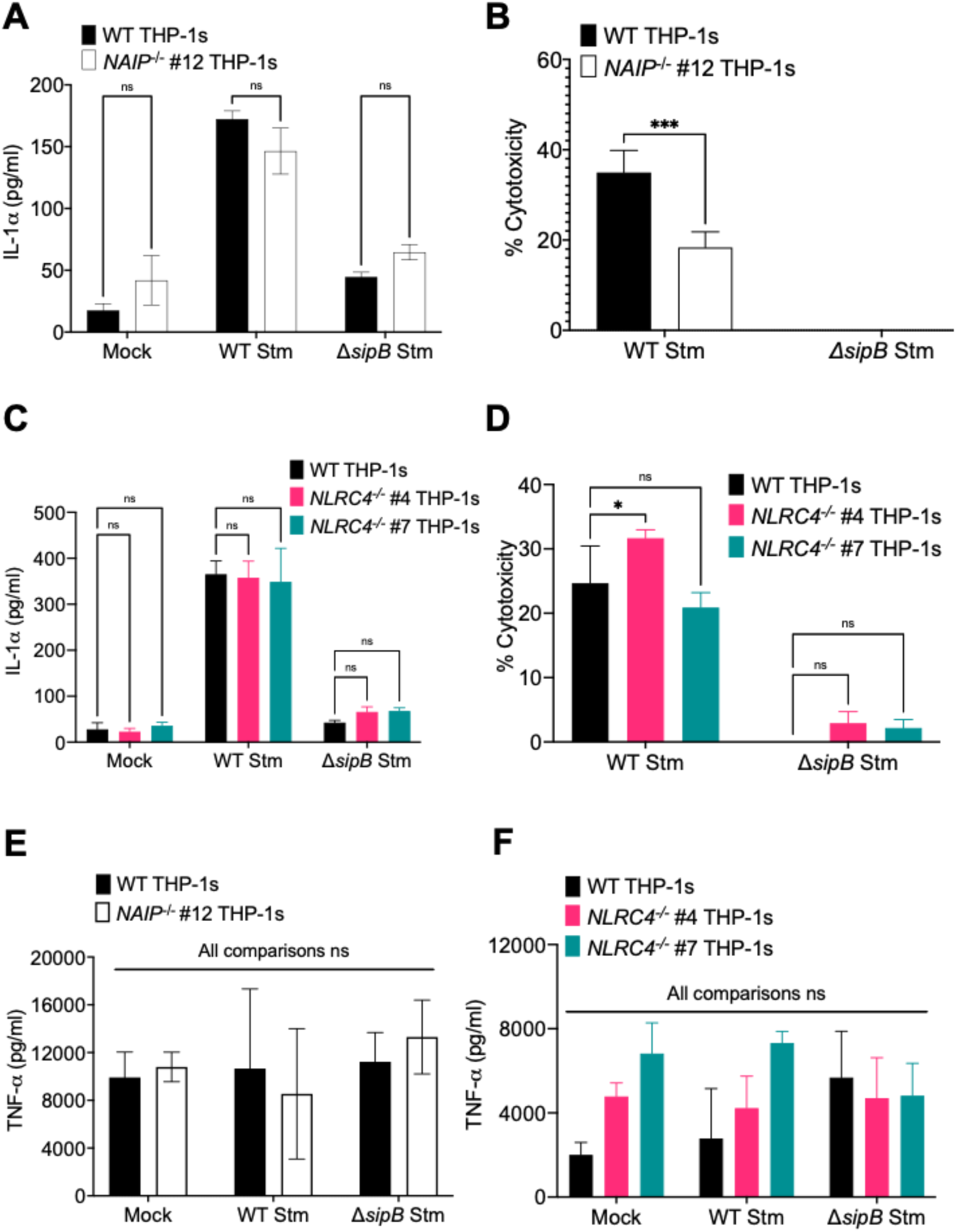
NAIP and NLRC4 are partially required for inflammasome activation during *Salmonella* infection in human macrophages. WT, *NAIP^-/-^,* or two independent clones of *NLRC4^-/-^* THP-1 monocyte-derived macrophages were primed with 100 ng/uL Pam3CSK4 for 16 hours. Cells were then infected with PBS (Mock), WT *S*. Typhimurium, or Δ*sipB S*. Typhimurium for 6 hours. As a control, cells were primed with 500 ng/mL LPS for 4 hours and treated with 10 uM nigericin for 6 hours. (A, C, E, F) Release of cytokines IL-1α and TNF-α into the supernatant were measured by ELISA. (B, D) Cell death (percentage cytotoxity) was measured by lactate dehydrogenase release assay and normalized to Mock-treated cells. ns – not significant, \**p* < 0.05, \*\*\**p* < 0.001 by Šídák’s multiple comparisons test (A, B, E) or by Dunnett’s multiple comparisons test (C, D, F). Data shown are representative of at least three independent experiments.

**S6 Fig. (related to Figure 3).**
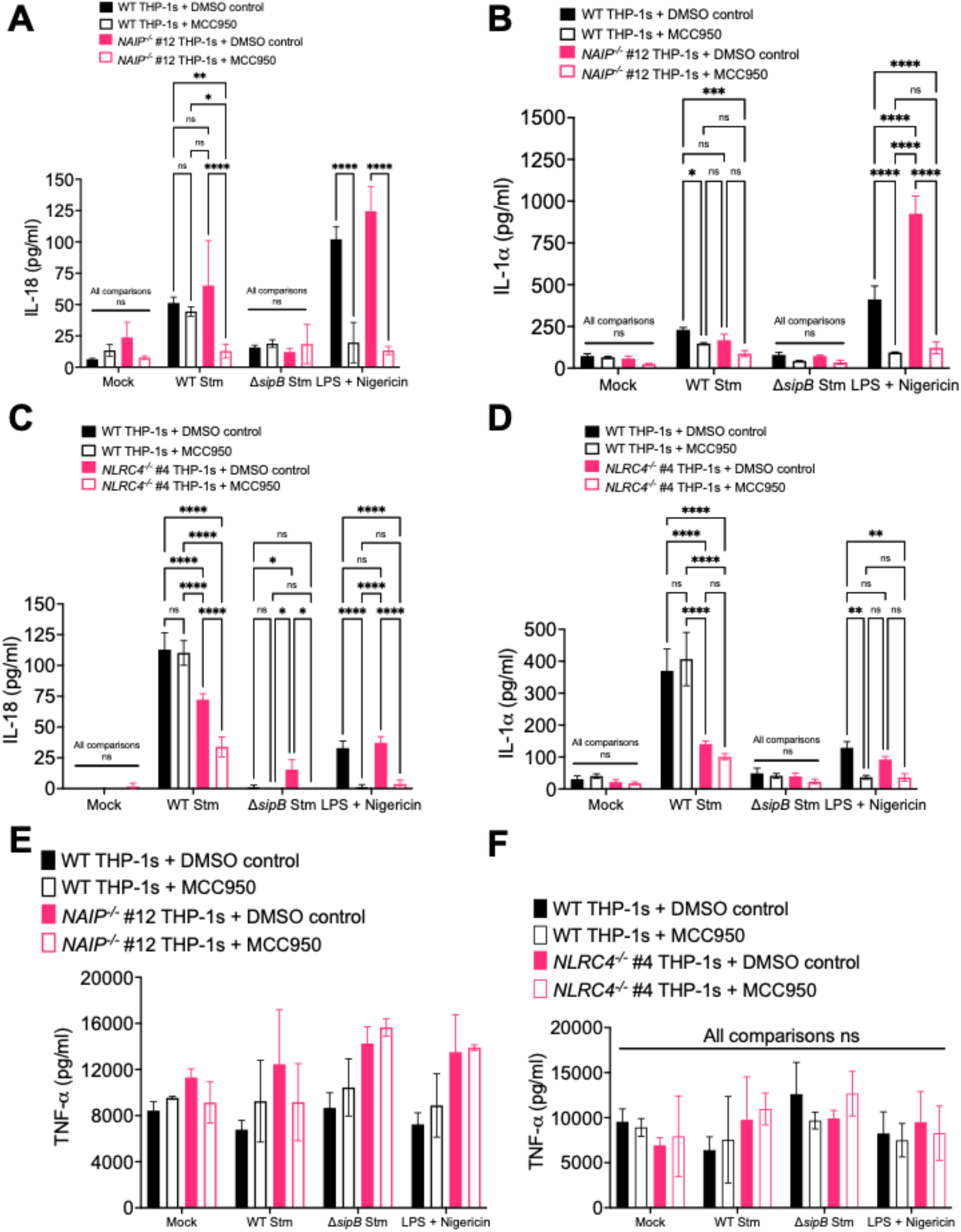
*Salmonella* induces NAIP/NLRC4- and NLRP3- dependent inflammasome activation in human macrophages. WT, *NAIP^-/-^,* or *NLRC4^-/-^* THP-1 monocyte-derived macrophages were primed with 100 ng/uL Pam3CSK4 for 16 hours. One hour prior to infection, cells were treated with 1 µM MCC950, a chemical inhibitor of the NLRP3 inflammasome. Cells were then infected with PBS (Mock), WT *S*. Typhimurium, or Δ*sipB S*. Typhimurium for 6 hours. (B) As a control, cells were primed with 500 ng/mL LPS for 4 hours and treated with 10 uM nigericin for 6 hours. (A-F) Release of cytokines IL-18, IL-1α, TNF-α into the supernatant were measured by ELISA. ns – not significant, \**p* < 0.05, \*\**p* < 0.01, \*\*\**p* < 0.001, \*\*\*\**p* < 0.0001 by Tukey’s multiple comparisons test.

**S7 Fig. (related to Figure 4).**
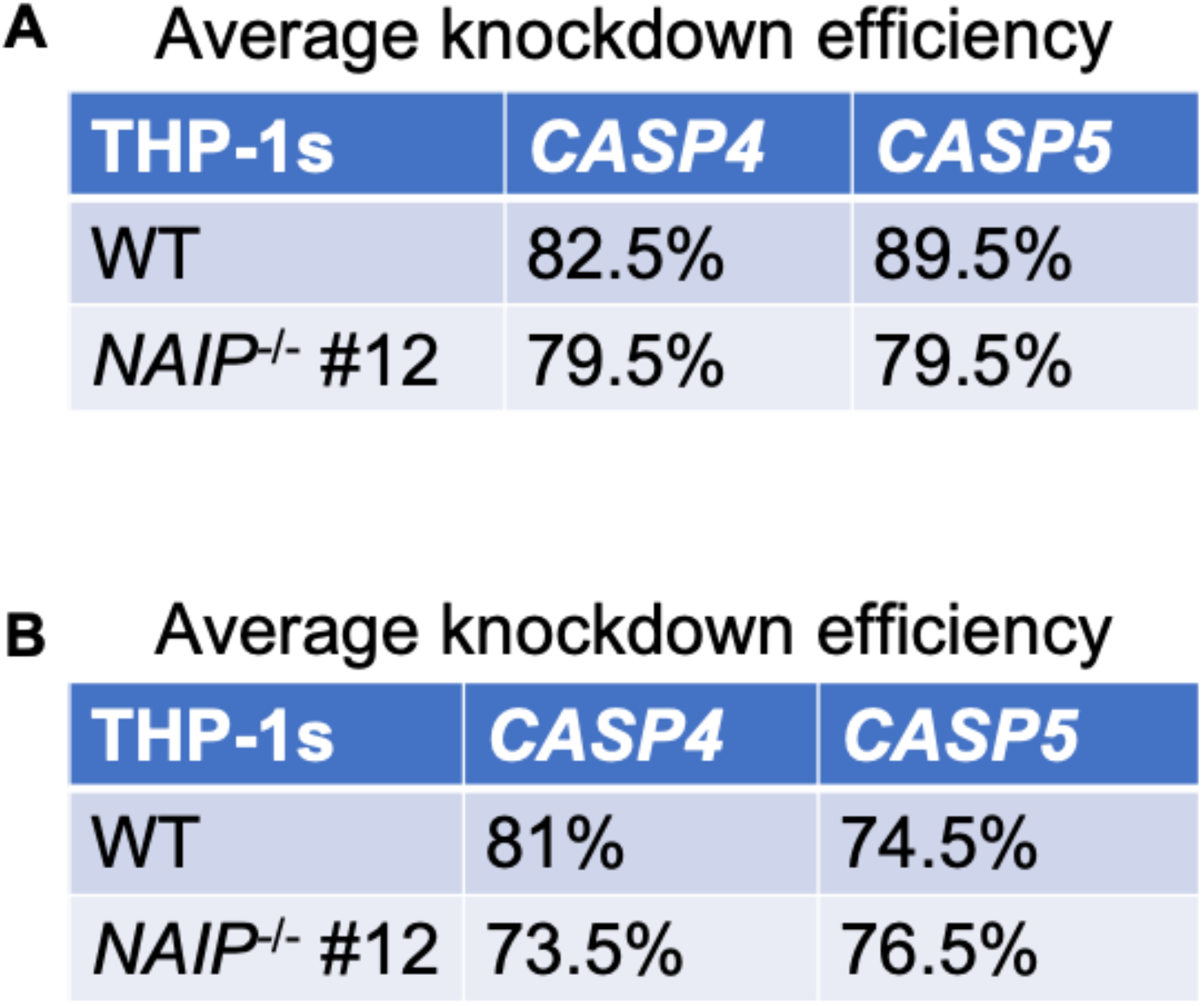
Knockdown efficiencies of siRNA-mediated silencing of *CASP4* and *CASP5* in human macrophages. Knockdown efficiencies following siRNA treatment were measured by qRT-PCR and normalized to housekeeping gene *HPRT*, and calculated relative to control-siRNA-treated cells. (A) siRNA targeting *CASP4* or *CASP5*. (B) siRNA targeting *CASP4* and *CASP5*. Data shown are averages of at least three independent experiments.

**S8 Fig. (related to Figure 5).**
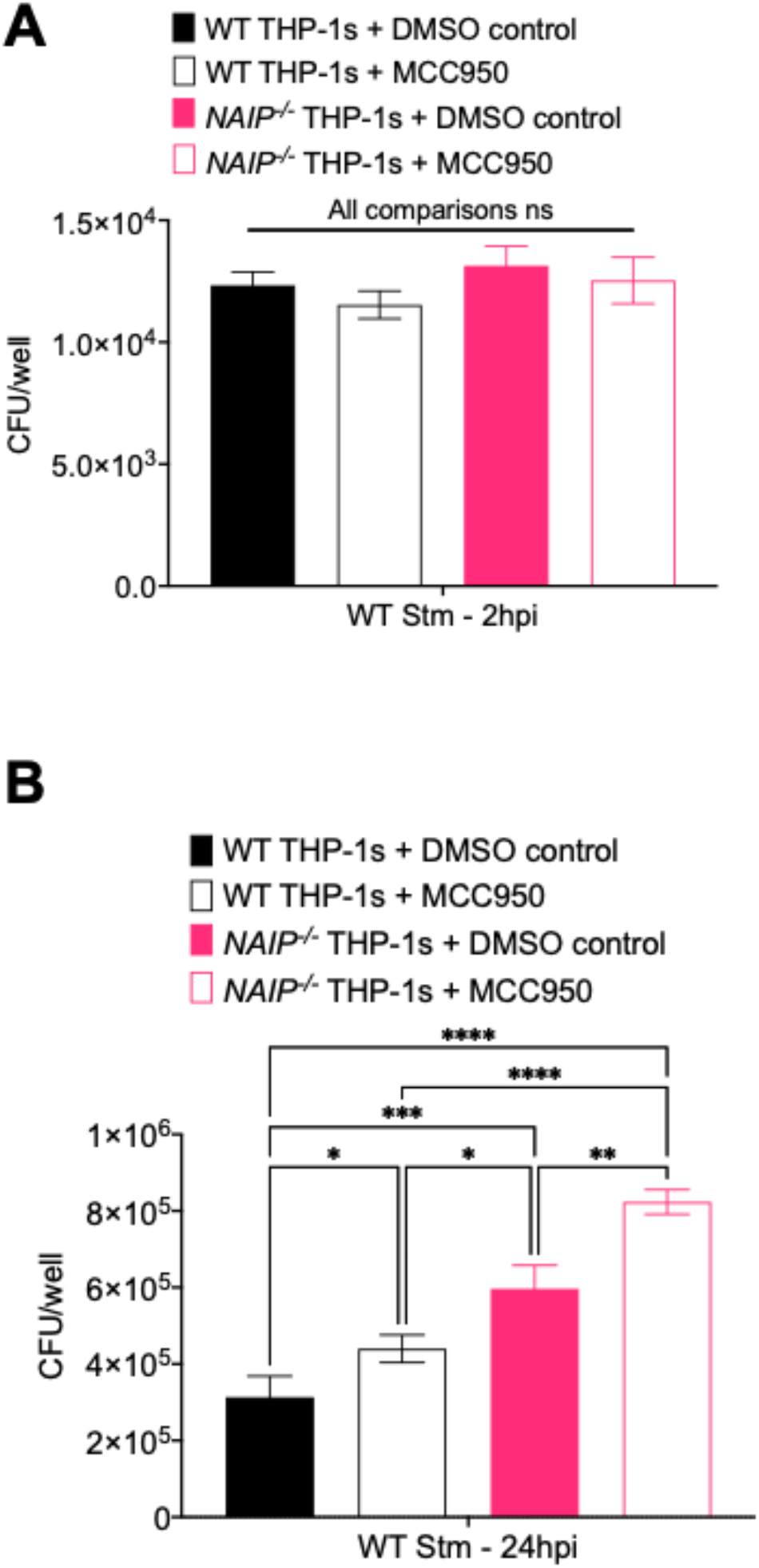
NAIP and NLRP3 restrict replication of *Salmonella* in human macrophages. WT or *NAIP^-/-^* THP-1 monocyte-derived macrophages were primed with 100 ng/ml Pam3CSK4 for 16 hours. One hour prior to infection, cells were treated with 1 µM MCC950 or DMSO as a control. Cells were then infected with WT *S*. Typhimurium. Cells were lysed at the indicated time points and bacterial were plated to calculate CFU. (A) CFU/well of bacteria at 2 hpi (B) CFU/well of bacteria at 24 hpi. \**p* < 0.05, \*\**p* < 0.01, \*\*\**p* < 0.001, \*\*\*\**p* < 0.0001 by Tukey’s multiple comparisons test. Data shown are representative of at least three independent experiments.

**S9 Fig. (related to Figure 6A).**
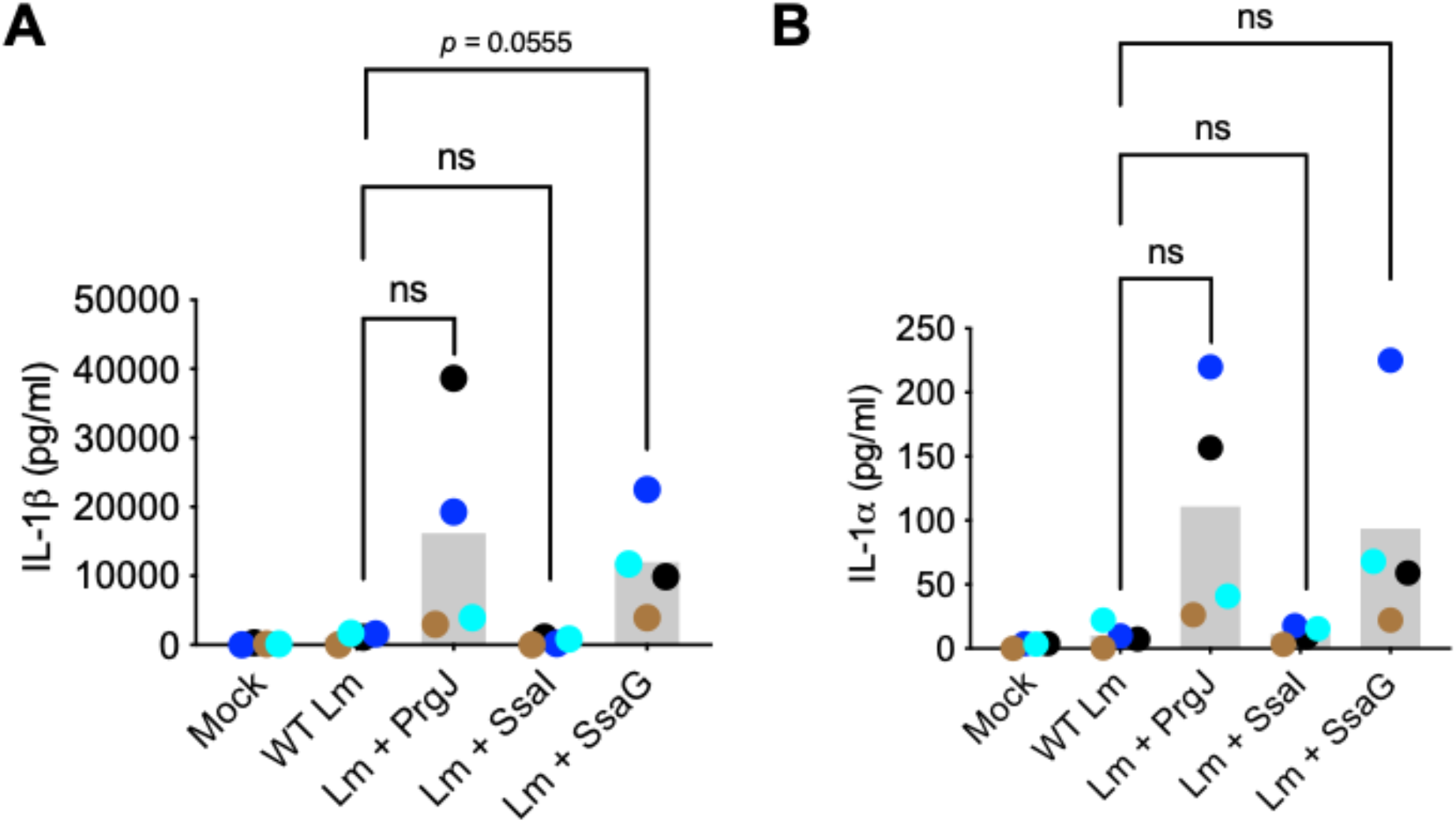
*Salmonella* SPI-2 needle protein SsaG activates the inflammasome in human macrophages. Primary hMDMs from four healthy human donors was infected with PBS (Mock), WT *Listeria* (WT Lm), *Listeria* expressing PrgJ (Lm + PrgJ), SsaI (Lm + SsaI), or SsaG (Lm + SsaG) for 16 hours at MOI=5. Each dot represents the triplicate mean of one donor. The grey bar represents the mean of all donors. Release of cytokines IL-1β and IL-1α, was measured by ELISA. *p* values based on paired t-tests.

**S10 Fig. (related to Figure 6B).**
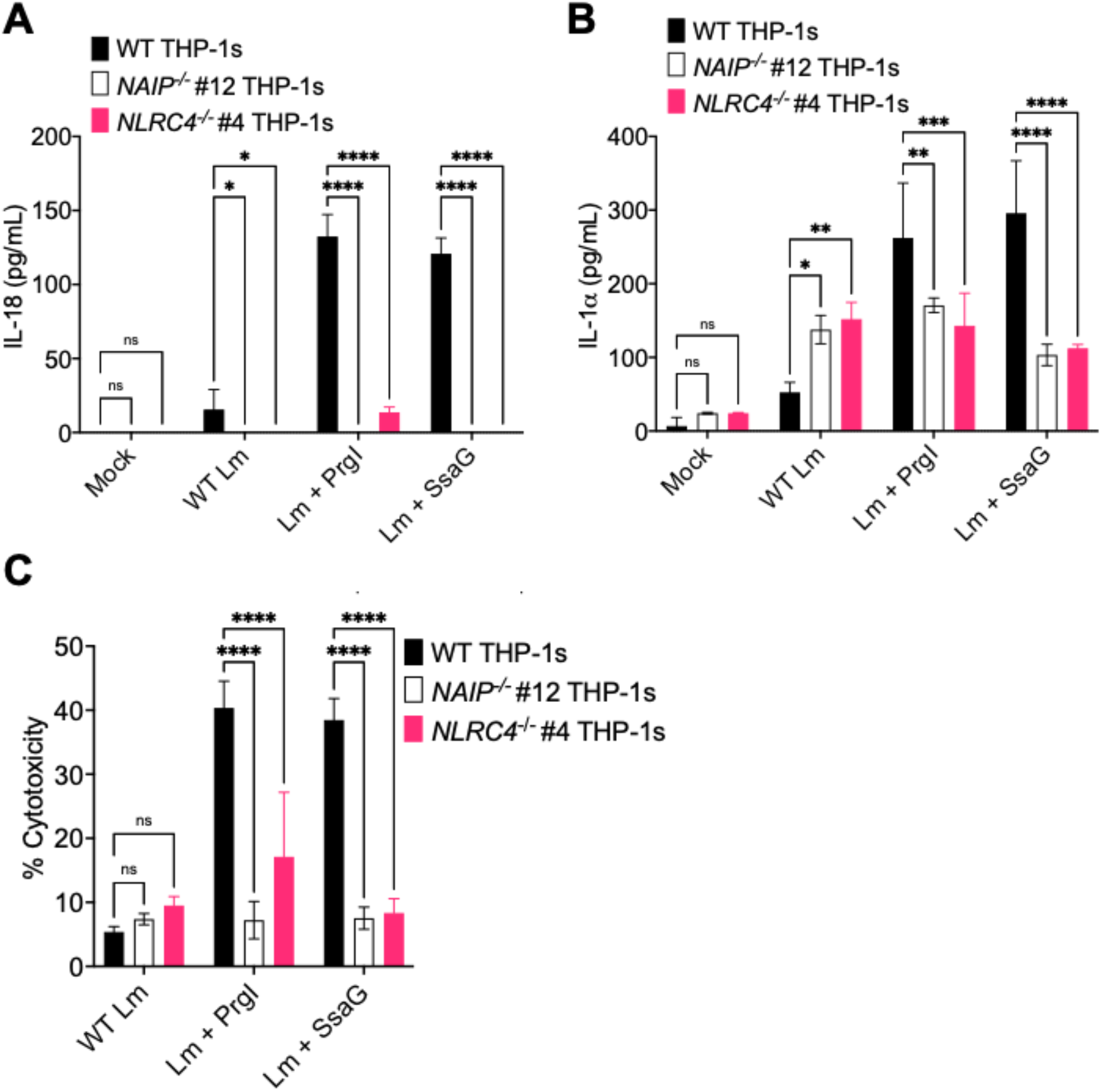
NAIP/NLRC4 are necessary for inflammasome responses to the *Salmonella* SPI-2 needle protein SsaG in human macrophages. WT, *NAIP^-/-^*, or *NLRC4^-/-^* THP-1 monocyte-derived macrophages were primed with 100 ng/ml Pam3CSK4 for 16 hours. Cells were then treated with PBS (Mock), WT *Listeria* (WT Lm), *Listeria* expressing PrgI (Lm + PrgJ) or SsaG (Lm + SsaG) for 6 hours at MOI=20. (A, B) Release of cytokines IL-18, and IL-1α was measured by ELISA. (C) Cell death was measured by lactate dehydrogenase (LDH) release. ns – not significant, \**p* < 0.05, \*\**p* < 0.01, \*\*\**p* < 0.001, \*\*\*\**p* < 0.0001 by Dunnett’s multiple comparisons test. Data shown are representative of at least three independent experiments.

**S11 Fig. (related to Figure 6C).**
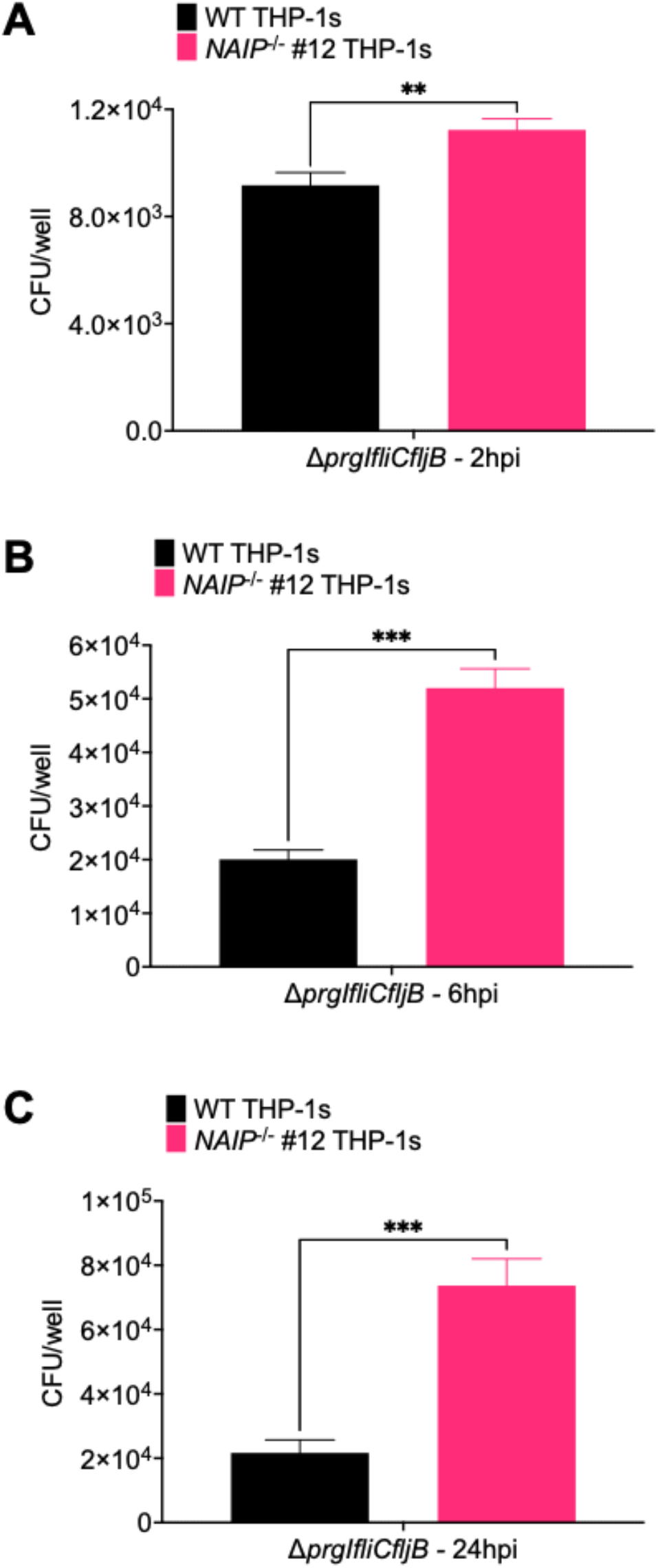
NAIP/NLRC4 inflammasome recognition of the SPI- 2 T3SS restricts *Salmonella* replication in human macrophages. WT or *NAIP^-/-^* THP-1 monocyte-derived macrophages were primed with 100 ng/ml Pam3CSK4 for 16 hours. Cells were then infected with a SPI-1 T3SS/flagellin-deficient strain of *S*. Typhimurium, Δ*prgIfliCfljB*. (A) CFU/well of bacteria at 2 hpi (B) CFU/well of bacteria at 6 hpi. (C) CFU/well of bacteria at 24 hpi. \*\**p* < 0.01, *\*\*\*p* < 0.001, by unpaired t-test. Data shown are representative of at least three independent experiments.

**S12 Fig.**
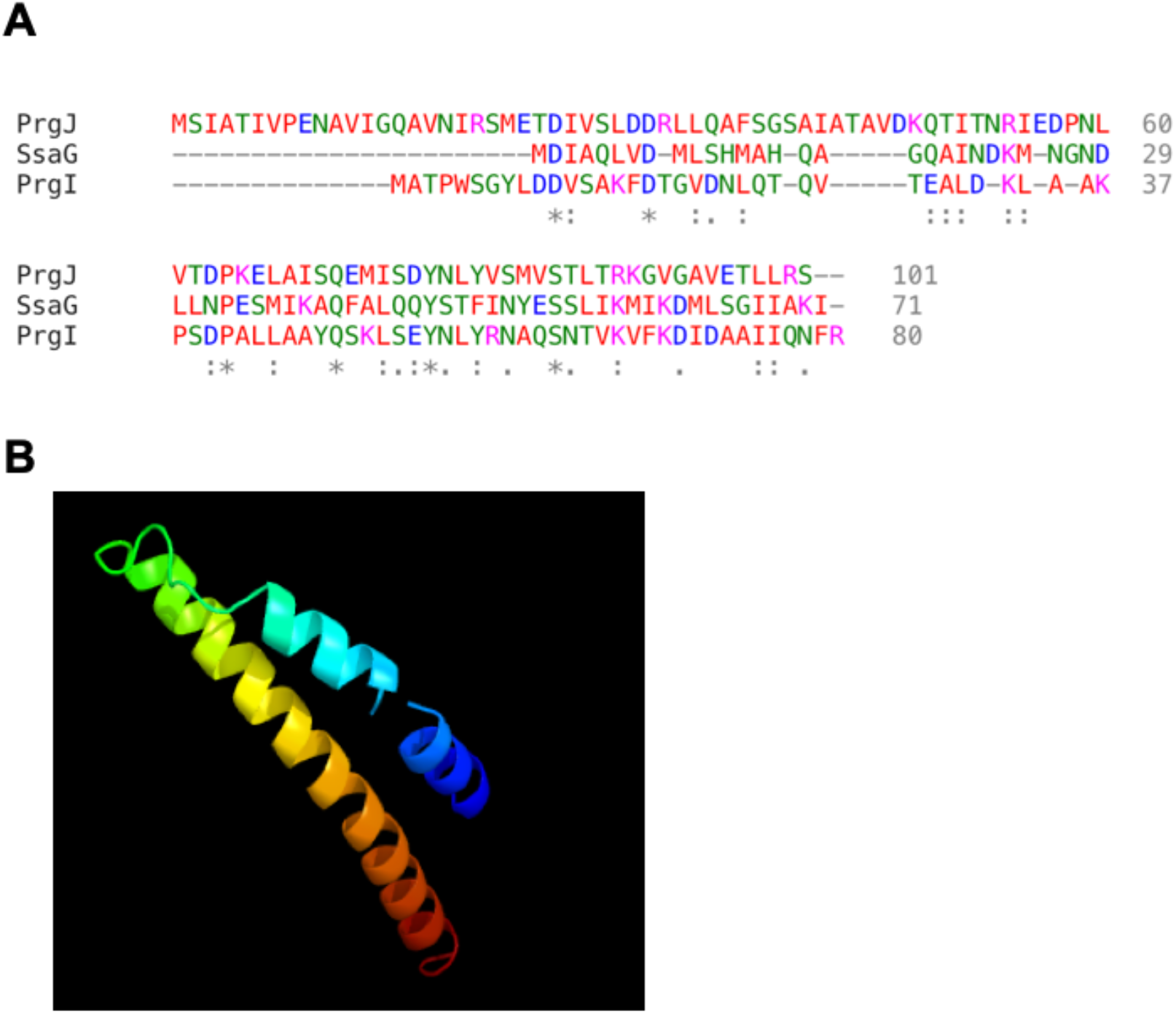
Sequence alignment and three-dimensional structural prediction of SsaG. (A) The primary sequences of PrgJ, PrgI, and SsaG were aligned using Multiple Sequence Alignment by Clustal Omega. **\*** indicates single, *fully conserved* residue, : indicates conservation between groups of *strongly* similar properties, and. indicates conservation between groups of *weakly* similar properties. Small, hydrophobic residues are indicated in red (AVFPMILW). Acidic residues are indicated in blue (DE). Basic residues are indicated in magenta (RK). The remaining residues are indicated in green (STYHCNGQ). (B) The three-dimensional structure of SsaG was predicted with high confidence and high coverage using the PHYRE2 server. The structure is colored from N to C terminus using the colors of the rainbow (red, orange, yellow, green, and blue).

## Notes

### Competing Interest Statement

The authors have declared no competing interest.

